# Effect of Terpenes from *Poria Cocos*: Verifying Modes of Action against Alzheimer’s disease Using Molecular Docking, Drug-induced Transcriptomes and Diffusion Network

**DOI:** 10.1101/2023.06.01.543358

**Authors:** Musun Park, Seo-Young Lee, Haeseung Lee, Jin-Mu Yi

**Author notes:** Correspondence should be addressed to: M.P. Korea Institute of Oriental Medicine, Daejeon, Republic of Korea.

## Abstract

*Poria cocos* is used to treat various diseases because of its unique terpenes. However, no study verifying its modes of action using only its compounds based on *in silico* experiments has been conducted. Here, molecular docking and drug-induced transcriptome analyses were performed to confirm the modes of action of *Poria cocos*. Additionally, a diffusion network analysis method was proposed to confirm its modes of action against Alzheimer’s. First, *Poria cocos* terpenes were collected from standard databases for molecular docking. They were then used for large-scale molecular docking using druggable proteomes, and the modes of action of lanostane and seco-lanostane, unique to *Poria cocos*, were confirmed. Additionally, the brain cell line was treated with *Poria cocos* to produce *Poria cocos*-induced transcriptome data, and the transcriptome-based modes of action of *Poria cocos* were confirmed. Finally, a diffusion network was constructed using Alzheimer’s, which acts on both modes of action, and submodules of Alzheimer’s with which terpenes interact were identified. It was confirmed that *Poria cocos* acts on the Wnt signaling pathway, Zn-to-anterograde axonal transport, autophagy impairment, insulin and AGE-RAGE signaling pathways, and apoptosis mechanisms. This study identified the modes of action of *Poria cocos* using biological data and *in silico* methods.

## Introduction

*Poria cocos* (PC), also known as Bok-ryeong, has long been used to treat various diseases. Because the terpenes contained in PC are known to have various pharmacological effects (1, 2), PC is highly valued as a therapeutic agent. PC has been used to treat edema caused by water retention in the body (3), and it shows various therapeutic effects when used together with other herbal medicines. For example, PC has a diuretic effect in Oryeong-san (4), antidiarrheal effect in Samryeongbaekchul-san (5), digestive function in Yukgunja-tang (6), and tranquilizing effect in Gwibi-tang (7). In addition, according to a recent study, PC is known to have various effects on blood pressure, diabetes, antitumor, anti-inflammation, immune regulation, and Alzheimer’s disease (AD) (8). Therefore, it is important to determine the modes of action (MOA) of PC as they can be used to treat various diseases.

The unique properties of PC differ from those of other herbal medicines. A unique feature of PC is that it belongs to the fungal family (9). In general, herbal medicines use plants, animals, and minerals, whereas fungi are rarely used. Mushrooms contain high amounts of lanostane and secolanostane terpenes; therefore, they may have different pharmacological mechanisms from those of other herbal medicines. (2). Previous studies have shown that drugs with similar scaffolds have similar mechanisms of action, demonstrating that the unique compounds in PC may have unique mechanisms (10, 11, 12). Therefore, studying PC-specific MOA can go beyond simply studying the MOA of herbal medicines and exploring new MOA that herbal medicines do not generally possess.

Because PC contains unique compounds, the MOA prediction results for the other herbal medicines were different. However, only a few studies have predicted the MOA of PC based on in silico experiments using PC compounds. Review articles on PC compounds and their MOA have been published (13, 14, 15). However, holistic MOA studies have not yet been conducted to elucidate the effects of PC. Therefore, it is necessary to study how the unique compounds of PC act on various biological MOA, such as diuresis, diarrhea, digestion, and tranquilization.

Several methods can be used to confirm the broad MOA of herbal medicines; however, molecular docking analysis (MDA) and drug-induced transcriptome analysis (DTA) are representative studies. MDA is a method for predicting whether a specific compound interacts with a specific protein by calculating the potential energy function and force field based on the molecular energy calculated from the structures of the compound and protein (16). DTA is a method of confirming drug response by measuring the relative expression level of mRNA in cells or specific tissues (17). If these methods were used, it would help elucidate the various MOA of PC.

MDA and DTA are excellent methods; however, they have obvious drawbacks. MDA is capable of confirming direct drug-target interactions, but not downstream drug actions. DTA is capable of confirming the transcription resulting from drug treatment, but not the upstream drug action. MDA and DTA have clear disadvantages, but because they are complementary to each other, it is expected that the complex MOA of herbal medicines can be revealed if the two methods are used together.

In this study, various biological MOA of PC were studied using the terpenes contained in PC. The upstream terpene mechanism, confirmed by drug-protein interactions, was confirmed through MDA. In addition, to identify the therapeutic effects of PC on neuropsychiatric and digestive diseases, DTA using brain and colon cell lines was performed to confirm the downstream mechanism of PC. Finally, an analytical method based on a diffusion network that connects the upstream and downstream mechanisms of PC was proposed (18), and the detailed pharmacological MOA of PC in AD pathology was confirmed.

## Results

### Terpene Collection from Poria Cocos

The terpenes included in PC were collected from several research papers and herb-compound databases and divided into six groups. Each group contained 123 lanostane-type triterpenes, 51 seco-lanostane-type triterpenes, six tricyclic diterpenes, five tricyclic diterpenes, 20 sterols, and 15 non-groups (Table 1, Supplementary Material 1). Among the collected terpenes, 125 druggable compounds with three-dimensional (3D) molecular structures were searched in the PubChem Database.

**Table 1.** Terpenes Collection Results of Poria Cocos

| Name | Reference | Dock-ability |
| --- | --- | --- |
| **Lanostane-type triterpenes** |  |  |
| Eburicoic acid | (51, 52, 53, 54, 55, 56, 57, 58, 59) | TRUE |
| O-acetylpachymic acid-25-ol | (60) | FALSE |
| 3β-Hydroxy-16α-acetoxy-lanosta-7,9(11),24-trien-21-oic acid | (56, 57, 60, 61, 62) | FALSE |
| Pachymic acid | (2, 45, 51, 52, 56, 59, 63, 64, 65, 66, 67, 68, 69, 70, 71, 72, 73, 74) | TRUE |
| lanostane | (75) | TRUE |
| Polyporenic acid C | (2, 51, 56, 59, 66, 67, 68, 69, 72, 74, 76, 77, 78, 79) | TRUE |
| 3-epi-Dehydrotumulosic acid | (56, 57, 58, 66, 68, 76, 80, 81, 82, 83) | TRUE |
| 25-Hydroxy-3-epidehydrotumulosic acid | (61, 77, 80, 84, 85) | TRUE |
| Versisponic acid E | (86) | TRUE |
| 3β,16α-Dihydroxylanosta-7,9(11),24-trien-21-oic acid | (3, 57, 58, 59, 61, 62, 68, 74, 82, 87, 88) | TRUE |
| Dehydroeburiconic acid | (52, 57, 77, 80, 84, 87, 89, 90) | TRUE |
| Daedaleanic acid B | (56, 61) | TRUE |
| 16α-Hydroxyeburiconic acid | (13, 52, 61, 77, 83) | TRUE |
| 3-epi-Dehydrotrametenolic acid | (52, 89, 91) | TRUE |
| Trametenolic acid | (53, 56, 57, 59, 72, 92, 93) | TRUE |
| Tumulosic acid | (3, 51, 55, 56, 59, 61, 65, 68, 69, 70, 83, 94, 95) | TRUE |
| Dehydrotumulosic acid | (2, 3, 45, 51, 56, 66, 68, 69, 72, 80, 95, 96, 97) | TRUE |
| 3-O-Acetyl-16α-hydroxytrametenolic acid | (13, 58, 61, 63, 66, 68, 80) | TRUE |
| 3-O-Acetyl-16α-hydroxydehydrotrametenolic acid | (57, 61, 62, 63, 80, 82, 93) | TRUE |
| 3-epi-Dehydropachymic acid | (56, 61, 62, 63, 66, 80) | TRUE |
| Dehydropachymic acid | (52, 56, 59, 63, 64, 65, 66, 68, 69, 73, 74, 82, 98, 99, 100) | TRUE |
| Dehydroeburicoic acid | (52, 53, 55, 57, 59, 61, 83, 93, 101) | TRUE |
| Dehydroeburicoic acid monoacetate | (102, 103, 104) | TRUE |
| 3β-p-Hydroxybenzoyldehydrotumulosic acid | (56, 57, 61, 62, 64, 72, 105) | TRUE |
| Dehydrotrametenolic acid | (2, 8, 45, 55, 56, 59, 64, 70, 74, 89, 93, 106, 107) | TRUE |
| Pinicolic acid A | (58, 86) | TRUE |
| masticadienoic acid | (8) | TRUE |
| 16α,27-Dihydroxydehydrotrametenolic acid | (61, 77) | TRUE |
| 15α-Hydroxydehydrotumulosic acid | (52, 61, 73) | TRUE |
| 5α,8α-Peroxydehydrotumulosic acid | (52, 61, 62) | TRUE |
| 16α,25-Dihydroxydehydroeburicoic acid | (52, 61, 77) | TRUE |
| Dehydrotrametenonic acid | (52, 57, 87) | TRUE |
| 25-Hydroxy-3-epitumulosic acid | (61, 77) | TRUE |
| Acetylburicoic acid | (56, 108) | TRUE |
| 16α,25-Dihydroxyeburiconic acid | (61, 77) | TRUE |
| 29-Hydroxydehydrotumulosic acid | (56, 61, 62, 69, 109) | TRUE |
| 3α-acetoxy-lanosta-8,24-dien-21-oic acid | (56) | FALSE |
| Ganoderic acid | (60) | TRUE |
| 6α-Hydroxypolyporenic acid C | (62, 67, 76) | TRUE |
| Pinicolic acid E | (56, 86) | TRUE |
| Eburicoic acid acetate | (104) | TRUE |
| 16α-Hydroxytrametenolic acid | (3, 13, 56, 57, 61, 62, 72) | TRUE |
| 3β,16α,30-Trihydroxy-24-methyl-lanosta-7,9(11),24(31)-trien-21-oic acid | (102) | TRUE |
| 16α-Hydroxy-3-oxo-24-methyl-lanosta-5,7,9(11),24(31)-tetraen-21-oic acid | (102) | TRUE |
| 3β,16α-Dihydroxy-7-oxo-24-methyl-lanosta-8,24(31)-dien-21-oic acid | (102) | TRUE |
| 3α,16α-Dihydroxy-7-oxo-24-methyl-lanosta-8,24(31)-dien-21-oic acid | (102) | TRUE |
| 29-Hydroxydehydropachymic acid | (56, 61, 69) | TRUE |
| 3-(2-Hydroxyacetoxy)-5α,8α-peroxydehydrotumulosic acid | (110) | TRUE |
| 3β-Acetoxylanosta-7,9(11),24-trien-21-oic acid | (56, 86, 102) | TRUE |
| 29-Hydroxypolyporenic acid C | (66, 67, 69) | TRUE |
| 3-Epi-pachymic acid | (61) | TRUE |
| 6,9-epoxyergosta-7,22-diene-3-ol | (95) | TRUE |
| 3β-Acetoxy-16α,24β-dihydroxylanosta-7,9(11),25-trien-21-oic acid | (102) | TRUE |
| masticadienolic acid | (8) | FALSE |
| Daedaleanic acid F | (86) | TRUE |
| Porilactone A | (86) | TRUE |
| Porilactone B | (86) | TRUE |
| Pinicolic acid F | (86) | TRUE |
| Ceanphytamic acid A | (111) | TRUE |
| Ceanphytamic acid B | (111) | TRUE |
| 15α-hydroxy-3-oxolanosta-8,24-dien-21-oic acid | (61) | FALSE |
| 16,27-dihydroxyl-dehydrotrametenolic acid | (56) | FALSE |
| 16-O-acetylpachymic acid | (56, 57, 58, 61) | FALSE |
| 16α-acetoxy-26,27-dimethoxyl-lanosta-8,24(31)-dien-21-oic acid | (58) | FALSE |
| 16α-acetoxy-lanosta-8,24-dien-21-oic acid | (58) | FALSE |
| 16α-acetoxypolyporenic acid C | (58) | FALSE |
| 16α-acetyloxy-24-methylene-3-oxolanosta-7,9(11)-dien-21-oic acid | (61) | FALSE |
| 16α-acetyloxyeburiconic acid | (61) | FALSE |
| 16α-Hydroxy-3-oxolanosta-7,9(11),24-trien-21-oic acid | (86) | FALSE |
| 16α-hydroxy-3-oxolanosta-8,24-dien-21-oic acid | (61) | FALSE |
| 16α-Hydroxydehydropachymic acid | (3) | FALSE |
| 16α-hydroxy-lanosta-7,9(11),24-trien-21-oic acid | (58) | FALSE |
| 16α-hydroxy-lanosta-8,24(31)-dien-21-oic acid | (58) | FALSE |
| 16α-hydroxy-lanosta-8,24-dien-21-oic acid | (58) | FALSE |
| 25-Hydroxy-3-epi-hydroxytumulosic acid | (80) | FALSE |
| 25-Hydroxypachymic acid | (56, 57, 58, 61, 67, 76, 77) | FALSE |
| 25-hydroxypolyporenic acid C | (58) | FALSE |
| 25α-hydroxytumulosic acid | (56) | FALSE |
| 3,15-O-diacetyl-dehydrotrametenolic acid | (112) | FALSE |
| 3,24-dioxo-16α-hydroxylanosta-7,9(11)-dien-21-oic acid | (56) | FALSE |
| 31-hydroxyl-16-O-acetylpachymic acid | (57) | FALSE |
| 3-acetyloxy-16α-hydroxytrametenolic acid | (56) | FALSE |
| 3-epi-(3'-hydroxy-3'-methylglutaryloxy)-16α-hydroxydehydrotrametenolic acid | (56) | FALSE |
| 3-epi-(3'-hydroxyl-3'-methylglutaryloxy)-dehydrotumulosic acid | (56) | FALSE |
| 3-epi-(3'-hydroxyl-3'-methylglutaryloxy)-tumulosic acid | (56) | FALSE |
| 3-epi-(3'-O-methyl malonyloxy)-dehydrotumulosic acid | (56) | FALSE |
| 3-O-acetyl-16α,26-dihydroxytrametenolic acid | (56, 72) | FALSE |
| 3-oxo-16α,25-dihydroxylanosta-7,9(11),24(31)-trien-21-oic acid | (62) | FALSE |
| 3-oxo-16α-hydroxylanosta-7,9(11),24(31)-trien-21-oic acid | (88) | FALSE |
| 3-oxo-16α-hydroxylanosta-7,9(11),24-trien-21-oic acid | (56) | FALSE |
| 3-oxo-6,16α-dihydroxylanosta-7,9(11),24(31)-trien-21-oic acid | (62) | FALSE |
| 3-oxo-6,16α-dihydroxytrametenolic acid | (62) | FALSE |
| 3α,16β-dihydroxylanosta-7,9(11),24-trien-21-oic acid | (56, 72, 113) | FALSE |
| 3β,5α,9α-trihydroxy-ergosta-7,22-diene-6-one | (95) | FALSE |
| 3β-acetoxy-16α-hydroxy-lanosta-8,24(31)-diene-21-oic acid | (88) | FALSE |
| 3β-Acetyloxy-16α-hydroxylanosta-7,9(11),24(31)-trien-21-oic acid | (98) | FALSE |
| 3β-Acetyloxy-16α-hydroxylanosta-7,9(11),24(31)-trien-21-oic acid | (98) | FALSE |
| 6α-hydroxydehydropachymic acid | (56, 57, 61, 62, 72) | FALSE |
| acetoxyeburiconic acid | (58) | FALSE |
| Coriacoic acid A | (114) | FALSE |
| Coriacoic acid B | (86, 114) | FALSE |
| Coriacoic acid C | (114) | FALSE |
| Coriacoic acid D | (114) | FALSE |
| Dehydrosulphurenic acid | (115) | FALSE |
| ergosta-4,22-diene-3-one | (95) | FALSE |
| ergosta-5,6-epoxy-7,22-dien-3-ol | (95) | FALSE |
| Hispidic acid B | (61) | FALSE |
| lanosta-7,9(11),24(31)-trien-21-oic acid | (58) | FALSE |
| lanosta-7,9(11),24-trien-21-oic acid | (58) | FALSE |
| lanosta-8,24(31)-dien-21-oic acid | (58) | FALSE |
| lanosta-8,24-dien-21-oic acid | (58) | FALSE |
| methyl trametenolate | (58) | FALSE |
| Methyl-O-acetylpachymate | (60) | FALSE |
| O-acetylpachymic acid | (60) | FALSE |
| Oxotrametenolic acid | (116) | FALSE |
| Pachymic acid methyl ester | (57, 60) | FALSE |
| Pinicolic acid | (56) | FALSE |
| Poriacosone A | (56, 57, 61, 68, 76) | FALSE |
| Poriacosone B | (56, 57, 61, 68, 76) | FALSE |
| Poricoic acid ZF | (117) | FALSE |
| Poricoic acid ZH | (117) | FALSE |
| Poricoic acid ZI | (118) | FALSE |
| Poricoic acid ZL | (118) | FALSE |
| **Seco-Lanostane-type triterpenes** |  |  |
| Poricoic acid A | (2, 55, 56, 59, 64, 74, 77, 84, 89, 106, 107, 109) | TRUE |
| Poricoic acid B | (2, 56, 59, 64, 72, 74, 77, 107, 109, 119) | TRUE |
| Poricoic acid G | (57, 58, 61, 68, 77, 85) | TRUE |
| Poricoic acid H | (61, 68, 77, 85, 120) | TRUE |
| Daedaleanic acid A | (86, 121) | TRUE |
| Poricoic acid E | (56, 80, 122, 123) | TRUE |
| Poricoic acid BM | (80, 83) | TRUE |
| Poricoic acid CM | (52, 57) | TRUE |
| 16-Deoxyporicoic acid B | (52, 57, 58, 61, 83, 91) | TRUE |
| 25-Hydroxyporicoic acid H | (52, 57) | TRUE |
| Poricoic acid D | (2, 52, 56, 58, 59, 61, 83, 105, 123) | TRUE |
| 26-Hydroxyporicoic acid DM | (56, 57, 77) | TRUE |
| 25-Hydroxyporicoic acid C | (56, 77, 83) | TRUE |
| 6,7-Dehydroporicoic acid H | (61, 77) | TRUE |
| Poricoic acid AM | (52, 56, 57, 58, 74, 77, 105, 106, 124) | TRUE |
| Poricoic acid DM | (52, 56, 59, 61, 77, 105) | TRUE |
| Poricoic acid GM | (56, 57, 61, 77) | TRUE |
| Poricoic acid HM | (56, 57, 61, 77, 83, 123) | TRUE |
| poricotriol A | (13) | TRUE |
| Poricoic acid C | (52, 56, 57, 58, 59, 61, 84, 105, 106, 123) | TRUE |
| Poricoic acid AE | (56, 57, 89) | TRUE |
| Poricoic acid CE | (56, 89, 123) | TRUE |
| 3-o-acetyl-dehydroeburicoic acid | (58) | TRUE |
| Daedaleanic acid D | (86) | TRUE |
| Poricoic acid I | (86) | TRUE |
| Poricoic acid J | (86) | TRUE |
| Poricoic acid K | (86) | TRUE |
| Poricoic acid L | (86) | TRUE |
| Poricoic acid M | (86) | TRUE |
| Daedaleanic acid E | (86) | TRUE |
| 11β-Ethoxydaedaleanic acid A | (111) | TRUE |
| 16-Deoxyporicoic acid BM | (86) | FALSE |
| 25-methoxy-29-hydroxyporicoic acid | (61) | FALSE |
| 26-hydroxyporicoic acid G | (56, 58, 123) | FALSE |
| 3,4-Secolanosta-4(28),7,9,24Z-tetraen-3,26-dioic acid | (91) | FALSE |
| poricoic acid GE | (56) | FALSE |
| poricoic acid HE | (56) | FALSE |
| Poricoic acid JM | (86) | FALSE |
| Poricoic acid N | (86) | FALSE |
| Poricoic acid O | (86) | FALSE |
| Poricoic acid ZA | (125) | FALSE |
| Poricoic acid ZC | (111) | FALSE |
| Poricoic acid ZD | (111) | FALSE |
| Poricoic acid ZG | (111, 117) | FALSE |
| Poricoic acid ZK | (86) | FALSE |
| Poricoic acid ZM | (111) | FALSE |
| Poricoic acid ZN | (111) | FALSE |
| Poricoic acid ZO | (111) | FALSE |
| Poricoic acid ZP | (111) | FALSE |
| Poricoic acid ZQ | (111) | FALSE |
| Poricoic acid ZR | (111) | FALSE |
| **Pentacyclic triterpenes** |  |  |
| Oleanolic acid | (59) | TRUE |
| hederagenin | (59) | TRUE |
| β-Amyrin acetate | (60) | TRUE |
| 3-O-acetyloleanolic acid | (126) | TRUE |
| oleanic acid 3-O-acetate | (95) | FALSE |
| triterpene acid | (127) | FALSE |
| **Tricyclic diterpenes** |  |  |
| (104)7-Oxocallitrisic acid | (104) | TRUE |
| Dehydroabietic acid | (104) | TRUE |
| Pimaric acid | (104) | TRUE |
| Dehydroabietic acid methyl ester | (128) | TRUE |
| 7-oxo-15-Hydroxydehydroabietic acid | (14) | TRUE |
| **Sterols** |  |  |
| Ergosta-7,22-dien-3-one | (126) | TRUE |
| β-Sitosterol | (95, 129) | TRUE |
| Ergosterol | (59, 107, 126) | TRUE |
| ergosta-7,22E-dien-3β-ol | (59) | TRUE |
| Ergosta-5,7-dien-3β-ol | (126) | TRUE |
| Ergosterol peroxide | (104) | TRUE |
| Daucosterol | (59, 129) | TRUE |
| (22E)-ergosta-5,7,9(11),22-tetraen-3β-ol | (126) | TRUE |
| Biemnasterol | (126) | TRUE |
| Cervisterol | (59, 95, 108) | TRUE |
| ergosterone | (130) | TRUE |
| 9,11-Dehydroergosterol peroxide | (104) | TRUE |
| Ergost-7-en-3β-ol | (126) | TRUE |
| Ergosta-4,22-dien-3-one | (126) | TRUE |
| 3β,5α-Dihydroxy-ergosta-7,22-dien-6-one | (95, 126) | TRUE |
| 6,9-Epoxy-ergosta-7,22-dien-3-ol | (126) | TRUE |
| (22E)-Ergosta-6,8(14),22-trien-3β-ol | (126) | FALSE |
| (22E)-Ergosta-8(14),22-dien-3β-ol | (126) | FALSE |
| (22E,24R)-ergosta-7,22-dien-3-one | (95) | FALSE |
| 3β,5α,9α-Trihydroxy-ergosta-7,-dien-6-one | (126) | FALSE |
| **Other compounds** |  |  |
| caprylic acid | (59) | TRUE |
| palmitic acid | (59) | TRUE |
| lauric acid | (59) | TRUE |
| mannitol | (95) | TRUE |
| ribitol | (95) | TRUE |
| Undekansaeure | (59) | TRUE |
| protocatechualdehyde | (93) | TRUE |
| (-)-pinoresinol | (93) | TRUE |
| Trimethyl citrate | (59) | TRUE |
| Ethyl glucoside | (59) | TRUE |
| 2-lauroleic acid | (59) | TRUE |
| Dimethyl L-malate | (59) | TRUE |
| L-uridine | (59) | TRUE |
| 6-hydroxykaempferol 7-glucopyranoside | (92) | FALSE |
| poricoic acid ZG | (117) | FALSE |

### Molecular Docking using Terpenes

#### Over-representation Analysis using Unique Terpenes of Poria Cocos

Twenty proteins were selected with docking probabilities greater than 50% in the lanostane and seco-lanostane groups and less than 50% in the non-lanostane groups (Table 2). Over-representation analysis (ORA) with 20 proteins predicted them to be effective in treating neurological and psychiatric diseases (Figure 3). Brain-related pathways, such as the Rap1 signaling pathway, nicotine addition, neuroactive ligand-receptor interaction, and cocaine addiction, were identified using KEGG (Figure 3A). Synapse-related actions and interleukin actions were confirmed using Gene Ontology biological processes (Figure 3B). It was also confirmed that it acts on AD and hypertension in OMIM (Figure 3C).

**Figure 1.**
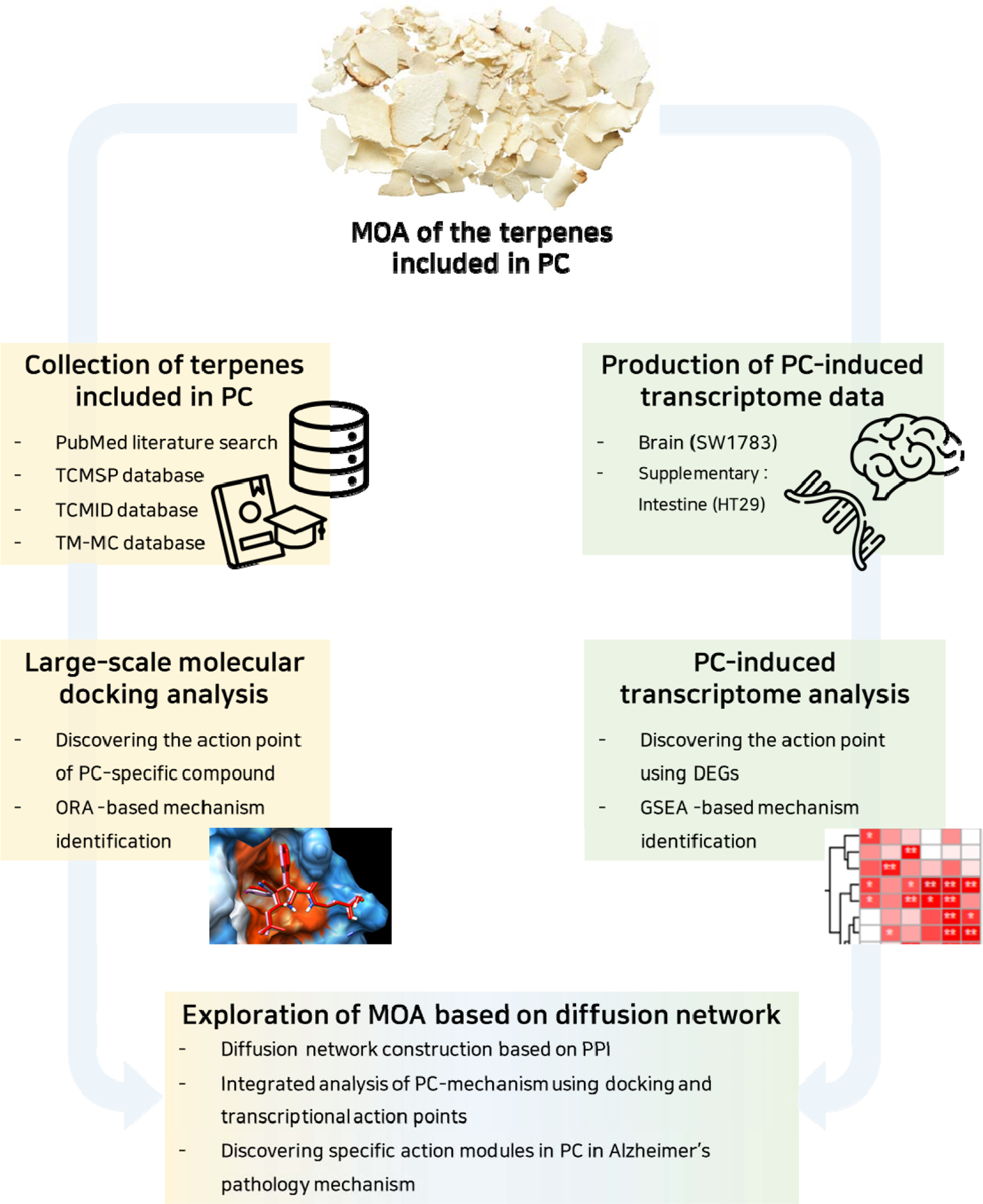
Framework for discovering the MOA of the terpenes included in PC Three methods were used to identify the MOA of terpenes included in PC. One is MDA method, which collects information on the terpenes contained in PC and finds druggable proteomes that binds to the terpenes. Another is DTA method, in which the MOA of PC was identified using the mRNA information obtained after processing PC into the cell line. Finally, DN that can be analyzed by integrating MDA and DTA results was constructed to confirm the detailed MOA of PC. MOA, Modes of action; PC, Poria Cocos; ORA, Over-representation analysis; DEGs, Differentially expressed genes, GSEA, Gene set enrichment analysis. MDA, Molecular docking analysis; DTA, Drug-induced transcriptome analysis; DN, Diffsion network.

**Figure 2.**
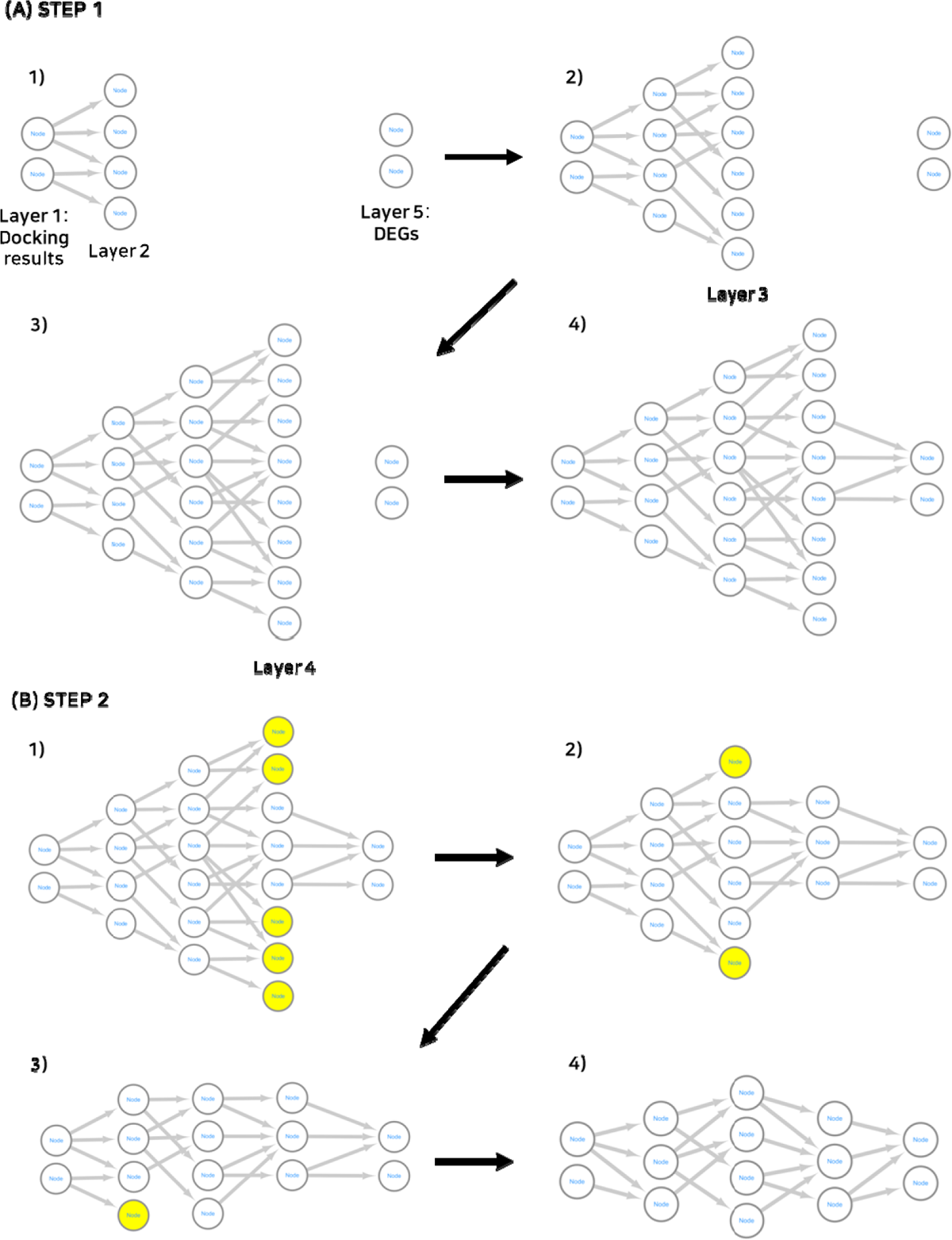
Method for construction of diffusion network The DN was constructed to integrate MDA results and DTA results. (A) DN construction method. Assign a new layer to the protein derived from the MDA results and the interacting proteins, and repeat the process of connecting the previous layer and the new layer three times to add a total of three layers. In the fourth layer, the network is completed by adding connections that interact with genes derived from DTA. (B) DN modification method. In the fourth layer, delete nodes that do not have a connection with the last layer. In this way, the network was completed by sequentially deleting nodes using the backpropagation method up to the second layer. DN, Diffusion network; MDA, Molecular docking analysis; DTA, Drug-induced transcriptome analysis.

**Figure 3.**
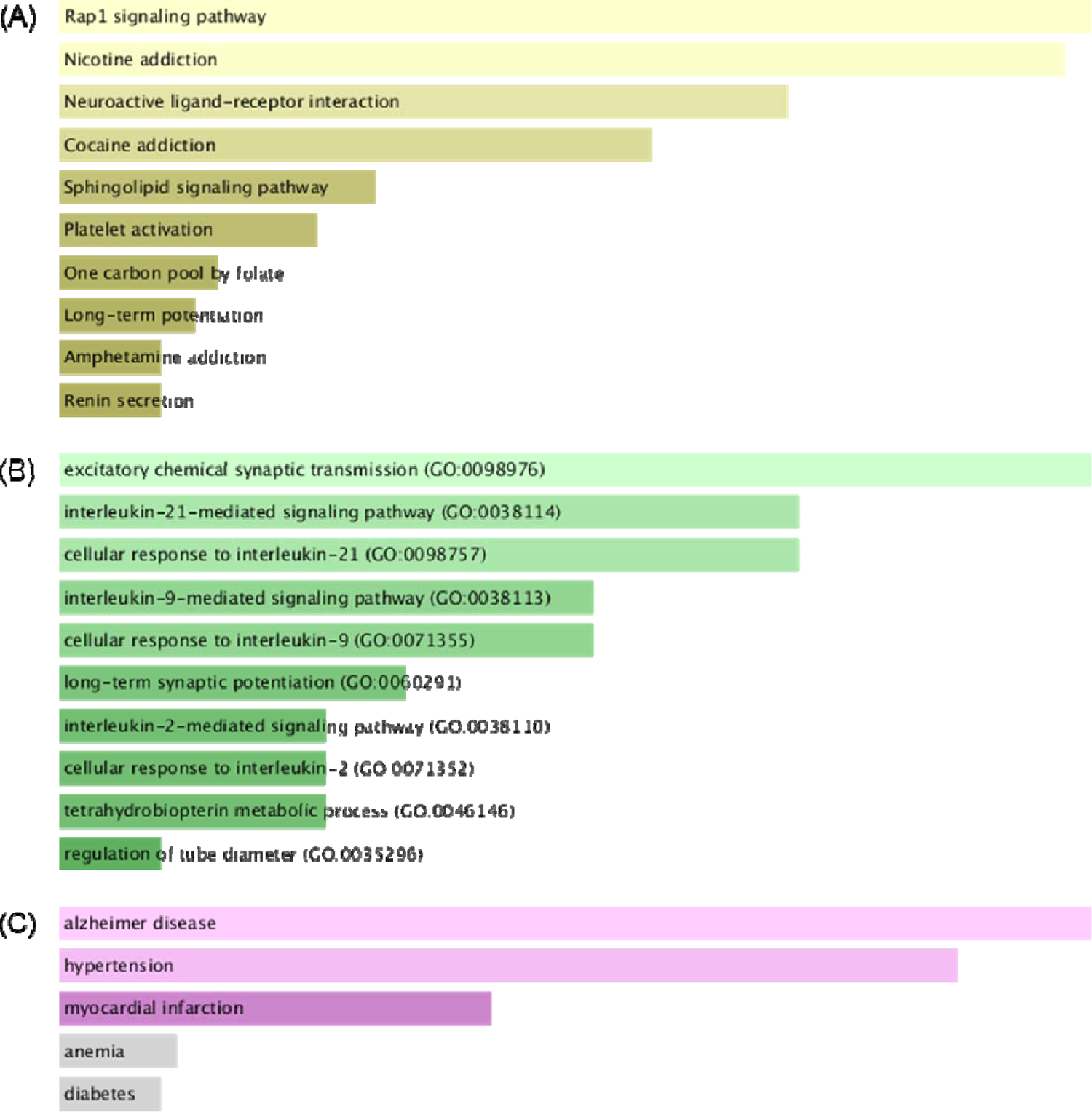
ORA results using lanostane terpenes and seco-lanostane terpenes ORA was performed on the EnrichR platform using proteins that specifically bind to lanostane and seco-lanostane terpenes, which are unique terpenes of PC. (A) ORA results using the KEGG gene set (B) ORA results using the GO biological process gene set. (C) ORA results using the OMIM disease gene set. ORA, Over-representation analysis. PC, Poria Cocos; KEGG, Kyoto Encyclopedia of Genes and Genomes; GO, Gene Ontology.

**Table 2.** Over-representation Analysis Results using Unique Terpenes of Poria Cocos

| Protein | Lanostane-type triterpenes | seco-Lanostane-type triterpenes | non-Lanostane-type |
| --- | --- | --- | --- |
| ITGAL | 0.910714 | 0.774194 | 0.263158 |
| ITGAX | 0.821429 | 0.806452 | 0.131579 |
| GRIN2B | 0.821429 | 0.903226 | 0.368421 |
| CHRM4 | 0.803571 | 0.967742 | 0.394737 |
| HCRTR1 | 0.785714 | 1 | 0.210526 |
| SLC6A8 | 0.785714 | 0.903226 | 0.394737 |
| SLC6A4 | 0.767857 | 0.709677 | 0.184211 |
| GRIN2A | 0.696429 | 0.967742 | 0.210526 |
| HCRTR2 | 0.678571 | 0.967742 | 0.263158 |
| PRKCZ | 0.678571 | 0.741935 | 0.236842 |
| PTGIS | 0.678571 | 0.645161 | 0.184211 |
| PRKCI | 0.660714 | 0.83871 | 0.315789 |
| DHFR | 0.642857 | 1 | 0.473684 |
| JAK3 | 0.642857 | 0.645161 | 0.394737 |
| JAK1 | 0.642857 | 0.935484 | 0.394737 |
| ACE | 0.625 | 0.677419 | 0.315789 |
| NOS3 | 0.607143 | 0.741935 | 0.447368 |
| SLC22A6 | 0.607143 | 0.709677 | 0.289474 |
| ADORA1 | 0.607143 | 0.935484 | 0.315789 |
| TFRC | 0.517857 | 0.870968 | 0.236842 |

#### Over-representation Analysis using Main Dockable Proteomes

As a result of the ORA using the main dockable proteomes, 54 pathways were found to be effective (Table 3, Supplementary Material 2). Fifty-four pathways showed various biological MOA possessed by PC. Biological terms such as the calcium signaling pathway, hypertrophic cardiomyopathy, and aldosterone synthesis and secretion are related to hypertension. Cortisol synthesis and secretion are associated with type II diabetes mellitus. Serotonergic synapses, GABAergic synapses, neurodegenerative pathways, and AD are biological terms related to neurological and psychiatric diseases.

**Table 3.** Top 25 prediction results of over-representation analysis results using main dockable proteomes

| KEGG Pathway | Docking count | P-value |
| --- | --- | --- |
| Calcium signaling pathway | 125 | 0.006 |
| cAMP signaling pathway | 123 | <0.001 |
| Cholinergic synapse | 115 | <0.001 |
| MAPK signaling pathway | 115 | <0.001 |
| Serotonergic synapse | 115 | <0.001 |
| Type II diabetes mellitus | 114 | <0.001 |
| Hypertrophic cardiomyopathy | 110 | <0.001 |
| Oxytocin signaling pathway | 107 | <0.001 |
| GnRH secretion | 102 | <0.001 |
| cGMP-PKG signaling pathway | 101 | <0.001 |
| GABAergic synapse | 100 | <0.001 |
| Adrenergic signaling in cardiomyocytes | 99 | <0.001 |
| Circadian entrainment | 98 | <0.001 |
| Arrhythmogenic right ventricular cardiomyopathy | 92 | <0.001 |
| Dilated cardiomyopathy | 91 | <0.001 |
| Cortisol synthesis and secretion | 90 | <0.001 |
| Aldosterone synthesis and secretion | 88 | <0.001 |
| Prion disease | 87 | <0.001 |
| PI3K-Akt signaling pathway | 84 | <0.001 |
| Pathways of neurodegeneration | 80 | <0.001 |
| Dopaminergic synapse | 79 | <0.001 |
| Renin secretion | 75 | <0.001 |
| Chemical carcinogenesis | 74 | <0.001 |
| Alzheimer disease | 73 | <0.001 |
| Cushing syndrome | 70 | <0.001 |

#### Gene Set Enrichment Analysis using Drug-induced Transcriptomes

GSEA based on the PC-induced transcriptomes of the SW1783 cell line revealed that many pathways were functional (Figure 4). We found that it was effective against neuropsychiatric diseases, such as AD, Parkinson’s disease, and Huntington’s disease, at low and medium concentrations. It acted as effectively as MDA in hypertrophic cardiomyopathy, but did not affect the calcium signaling, aldosterone-related, and diabetes-related pathways. In the hypertrophic cardiomyopathy pathway, it acts as effectively as MDA. However, it did not affect the calcium signaling pathway, aldosterone-related pathway, or diabetes-related pathway, showing different results from those of MDA. In addition, it was confirmed that PC was effective against bacterial infections by examining pathways such as *Vibrio cholerae* infection, pathogenic *Escherichia coli* infection, and viral myocarditis. GSEA of HT29 cells revealed many biological terms related to metabolism (Supplementary Figure 1). Similar to the results for SW1783, it was confirmed that it also acts against *Vibrio cholerae* infection.

**Figure 4.**
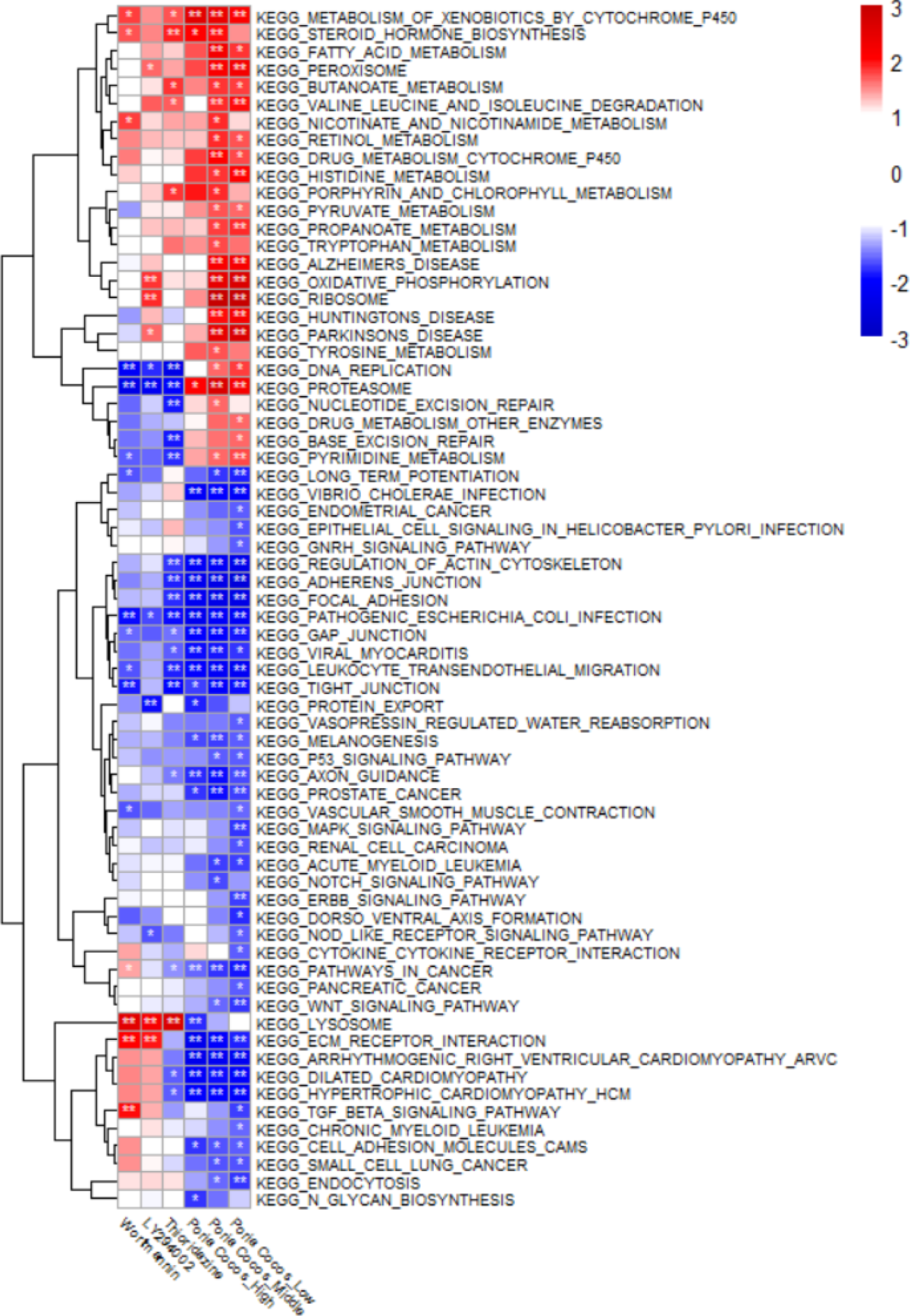
GSEA results using the SW1783 cell line-based PC-induced transcriptomes GSEA was performed using the KEGG pathway gene set. Poria Cocos_High, Poria Cocos_Middle, and Poria Cocos_Low are the results of analyzing the gene expression values obtained by treating the PC water extract with high (500 ug/mL), medium (100 ug/mL), and low (20 ug/mL) concentrations, respectively. Wortmannin, LY294002, and Thioridazine are positive controls. Wortmannin and LY294002 are known to inhibit inflammation by acting on the cell cycle mechanism, and Thioridazine is known to dopaminergic receptor antagosist. GSEA, Gene set enrichment analysis. PC, Poria Cocos; KEGG, Kyoto Encyclopedia of Genes and

### Diffusion Network Analysis (Figure 5)

#### Alzheimer’s Diffusion Network Construction (*Figure 5A*)

Diffusion network construction began with 14 proteins (layer1) from the MDA results and 22 proteins (layer5) from DEGs. Network construction identified 84, 169, and 24 proteins in layers 2, 3, and 4, respectively. Finally, only nine proteins remained in layer 1 and eight proteins remained in layer 5.

**Figure 5.**
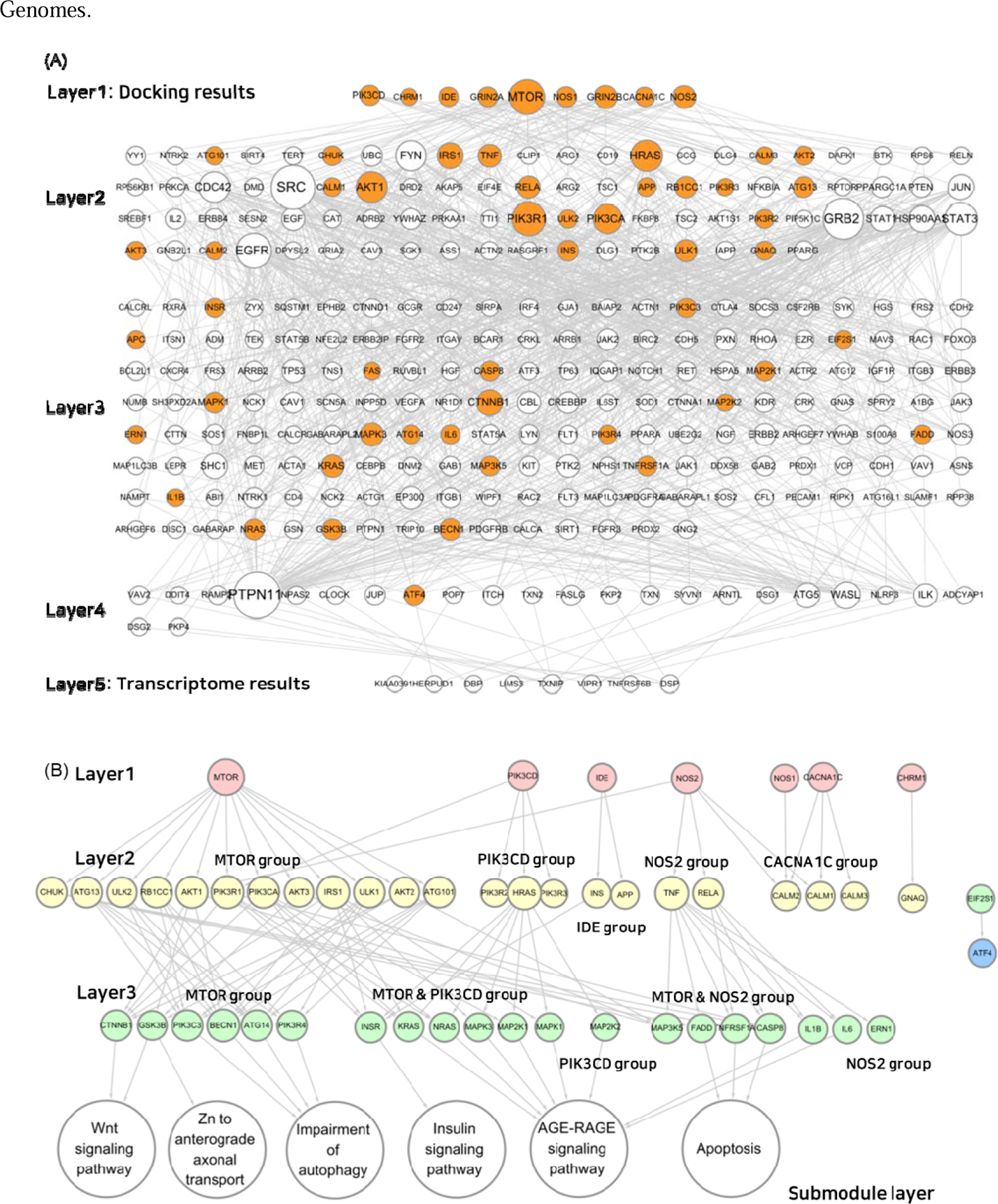
MOA analysis result of PC using AD-related DN (A) Diffusion network associated with AD. In layer 1, terms related to AD were selected from the results of MDA, and in layer 5, DEGs were selected and a diffusion network was built using them. Orange circles indicate proteins included in the KEGG AD gene set. (B) MOA analysis result of PC using AD-related DN. Each layer was rebuilt using only the orange circles in the network above. Layers 2 and 3 were grouped so that it was possible to identify which protein signal transduction was initiated in layer 1. It was confirmed which AD-submodule each group specifically acts on using KEGG mapper. MOA, Modes of action; PC, Poria Cocos; AD, Alzheimer’s disease; DN, Diffusion network; MDA, Molcular docking analysis, DEGs, Differentially expressed genes; KEGG, Kyoto Encyclopedia of Genes and Genomes.

#### Submodule Construction for Identifying Modes of Action

Proteins belonging to the Kegg Alzheimer’s pathway gene set (KAGS) were color-mapped to the Alzheimer’s diffusion network (ADN) (Figure 5A). In Layer 1, all nine genes were included in the gene set (9/9). The deeper the layer, the lower is the probability of belonging to a gene set. Only 23 proteins (23/84) in layer 2, 23 proteins (23/169) in layer 3, and 1 protein (1/24) in layer 4 belonged to the gene set. In the last DEG layer, the gene set did not contain any proteins belonging to the KAGS (0/8).

Only proteins belonging to the KAGS were selected separately and the network was reconstructed to confirm which submodules in layer 3 were associated (Figure 5B, Supplementary Figure 2). As a result of this analysis, seven proteins were selected from layer 1 except for two without connections. In layer 2, we rearranged the groups interacting with layer 1 into MTOR, PIK3CD, IDE, NOS2, and CACNA1C groups. Layer 3 was also rearranged into MTOR, MTOR, PIK3CD, PIK3CD, MTOR, NOS2, and NOS2 groups based on major connectivity. Each group was linked to related AD submodules. The MTOR group was related to the Wnt signaling pathway and Zn to anterior axonal transport and autophagy impairment mechanisms (Supplementary Figures 2A and 2B). The MTOR and PIK3CD groups acted on the insulin signaling pathway (Supplementary Figure 2C), whereas the PIK3CD group acted on the AGE-RAGE signaling pathway (Supplementary Figure 2D). The MTOR and NOS2 groups were confirmed to be associated with apoptosis (Supplementary Figure 2E).

## Discussion

In this study, in silico MDA, *in vitro* DTA, and DN analyses were conducted to identify the various mechanisms of PC using PC-terpenes. In clinical medicine, PC is known to be effective against various conditions, such as high blood pressure, bacterial infections, decreased metabolism, and mental disorders. In this study, to investigate the therapeutic effect of PC in clinical medicine, the pharmacological MOA of PC was analyzed using various techniques, and PC was confirmed to be particularly effective in AD.

This study had four important implications. First, the therapeutic effects of lanostane-type terpenes and seco-lanostane-type terpenes, which are unique compounds of PC, were identified using MDA. Second, a new mathematical model-based analysis method that can utilize even small amounts of compounds contained in PC by analyzing dockable proteomes was proposed. Third, the clinical efficacy of PC and the results of previous studies were verified using in silico MDA and *in vitro* DTA. Finally, to study the MOA of herbal medicines, the relationship between MDA was identified to confirm the upstream MOA and DTA, and a DN-based analysis method for integrated analysis was proposed.

PC contains unique compounds, including lanostane and seco-lanostane terpenes. Terpenes are unsaturated hydrocarbons produced from natural products and are known to have antioxidant, anti-inflammatory, and anticancer effects (41). Terpenes are also included in other herbal medicines, and because they have a different scaffold structure than lanostane-based terpenes, they interact with different proteins. This approach is the basis for explaining the physiological activity of PC compared to that of other medicinal herbs. For example, triterpenes with five rings, such as oleanolic acid and hederagenin, can act on pathways involving cholinergic synapses, type 2 diabetes, and GnRH secretion (12). In comparison, lanostane triterpenes most likely acts on the Rap1 signaling pathway, nicotine addiction, neuroactive ligand-receptor, and cocaine addiction pathways (Figure 3A). The Rap1 signaling pathway is associated with the blood-brain barrier (42), and all other pathways are related to neuropsychiatric disorders. In addition, interleukin-21 derived from the biological process of lanostane triterpenes, is already well known to be associated with AD (43) (Figure 3B). Therefore, the fact that PC is effective against mental illnesses is a unique characteristic of PC. Because PC has a high possibility of pharmacological MOA, which animal and plant herbal medicines cannot, research on PC will become more important in the future.

There is a problem with multi-compound-based pharmacological mechanism studies using herbs that do not consider the amount of compounds. Owing to the characteristics of herbal medicines, it is very difficult to analyze the pharmacological MOA of herbs by considering the amounts of compounds because the compound content varies depending on the cultivation area and harvesting time. To overcome the limitations of quantitative analysis, several compounds included in PC were analyzed, and the results were interpreted based on the number of pathway appearances. (Table 3). This approach is based on the assumption that an effect emerges when several compounds are combined. This is based on 1) the fact that compounds with the same scaffold structure have similar pharmacological mechanisms (11, 12), and 2) that multiple ligand docking analysis is possible even with different structures (44). However, because this approach considers each pathway to be acting on multiple compounds, it may act on a wider range of pathways than presently known. To overcome these drawbacks, a permutation test was performed. This statistical approach was used to limit the analysis of the MOA of the PC, which could be overanalyzed. The new mathematical model-based drug mechanism prediction method proposed in this study is significant because it indirectly estimates and uses the amounts of compounds that are not considered in multi-component herbal medicine studies. From a clinical perspective, the efficacy of PC against high blood pressure, diabetes, and mental disorders has been well predicted. However, as the exact amount was not considered, in future studies, MOA studies considering the amounts of herbal compounds should be conducted to clearly reveal the MOA of herbal medicine.

One of the characteristics of this study is that the pharmacological MOA upstream and downstream of PC were significantly different. MOA for conditions such as high blood pressure, diabetes, and psychiatric disorders were confirmed using MDA, and MOA for conditions such as psychiatric disorders, bacterial infections, and metabolism were identified using DTA. In this study, the MOA of MDA and DTA were analyzed differently; however, the results of both analyses revealed effective pharmacological MOAs for PC as in previous studies on diabetes, immune responses, and lipid metabolism. PC was shown to have anti-diabetic effect in diabetic mice through a mechanism that involved glucose-lowering activity and sensitizing PPAR-gamma independent insulin-mediated glucose uptake (45). PC also regulates immune responses by increasing NK cell activity and INF-γ secretion and decreasing IL-4 and −5 secretion in mouse splenic T lymphocytes (46). A previous study reported that PC is also effective in reducing serum cholesterol and triglyceride levels in high-fat-diet model, and regulates lipid homeostasis and bile acid metabolism via FXR/PPARα-SREBPs signaling pathway (47). Although the results from the two types of analysis were very different, the pharmacological MOAs for psychiatric disorders were consistent. In a previous study, PC was shown to be effective in treating psychiatric disorders. In studies to alleviate or modulate these symptoms, such as apathy, depression, sleep disruption and psychosis, which are the core features of patients with AD, Liu et al. identified that the extract of PC can ameliorate cognitive function by reducing Amyloid-β formation and improving Ab clearance in AD Mouse model (48). In addition, a study has shown that PC can regulate sleep architecture via GABA_A_-ergic mechanism, which increases the protein expression level of GAD65/67 and GABA_A_ receptors subunits in primary hypothalamic neuronal cells (49). Taken together, the pharmacological MOAs derived from these two analyses can explain not only the herbal efficacy of PC but also the latest research findings. Both MDA and DTA have advantages and disadvantages; however, because they are complementary to each other, it would help to use both assays together in future MOA studies.

The need for a method that can integrate and explain the results of psychiatric disorders, that were found to be equally effective in two analyses with different results, was confirmed, and a new analysis method based on a diffusion network was introduced (Figure 2). A diffusion network was constructed using the AD pathway, which confirmed that both MDA and DTA had significant effects, and the pathological MOA was successfully identified (Figure 5, Supplementary Figure 2). Two important findings were derived from this analysis. First, the AD pathway provided by KEGG is composed of a large number of biological terms and subnetworks, and specific subnetworks in which PC acts can be identified using the diffusion network. Second, terms related to the KEGG AD pathway gradually disappeared as the diffusion network layer deepened; therefore, the disease-related KEGG pathway did not fully reflect the results of the transcriptome analysis (Figure 5A). The KEGG pathway is a very useful database containing many biological terms such as genes, proteins, RNAs, compounds, glycans, chemical reactions, disease genes and other biological entities, but it is too complex and extensive. It is estimated that it will be difficult to include all results of transcriptome analysis. The model proposed in this study can reduce and model complex physiological and pharmacological mechanisms and identify information not provided by existing databases; therefore, it can be used in drug research.

This study has some limitations. The polysaccharides present in PC (50) could not be subjected to molecular docking analysis because their molecular weights were too large. In DTA, it is assumed that polysaccharides with high molecular weights will be filtered out during the sterilization process of the extract; therefore, there is no problem with our research strategy of using only terpenes. Although the effective function of polysaccharides is still being studied (50), it is difficult to perform *in silico* analyses to predict the pharmacokinetic and pharmacodynamic properties of polysaccharides. Therefore, techniques to predict the pharmacological MOA of compounds with very large molecular weights, such as polysaccharides, are needed in the future.

In the present study, the therapeutic efficacy of PC was verified using *in silico* MDA and *in vitro* DTA assays. In particular, the MOA of PC, which is widely used in clinical medicine, was identified using various biological data.

In summary, the pharmacological MOA of terpenes in PC were identified. To identify extensive pharmacological MOA of PC-terpenes, *in silico* MDA was performed by collecting PC-terpenes from research papers and herbal compound databases. In addition, *in vitro* DTA using the PC extract was performed to confirm the pharmacological MOA, and the MDA and DTA results were integrated using the diffusion network. As a result of the analysis, although the MDA results and the DTA results did not match, it was confirmed that both results were included in the clinical efficacy of PC. To integrate the two different results, a diffusion network was used for the analysis. Thus, the pharmacological MOA of PC-terpenes acting from upstream to downstream were identified. This study used research methods that have not been used in previous studies. Based on the molecular scaffold-based quantitative structure-activity relationship theory, we presented an analytical model to predict biological MOAs by combining small amounts of compounds with other compounds. We also presented a diffusion network, a research model that can confirm the diffusion of drug action mechanisms by linking the upstream and downstream regions of the pharmacological MOA. The method used in this study is expected to be useful for identifying pharmacological MOA based on complex natural products.

## Materials and Methods

### Overview of the Study

In this study, we investigated the biological MOA of PC using the terpenes present in PC. First, compound information was collected from the PubMed and herbal compound databases. To confirm the MOAs of PC upstream, MDA was performed using human druggable proteomes and the collected terpenes. Next, to confirm the MOA of PC downstream, DTA was performed using the HT29 and SW1783 cell lines. Finally, using the AD analyzed as the MOA point that works in both the MDA and DTA results, an AD-based diffusion network linking the results from both the analyses was established to identify the detailed pharmacological MOA of PC (Figure 1).

### *In silico* Molecular Docking Analysis

#### Method for Collecting Terpenes Contained in Poria Cocos

Various sources were used to collect as much PC terpene information as possible. First, the terpenes included in 23 papers found by searching for “*Poria Cocos*” and “HPLC” in PubMed database were collected. In addition, terpenes not found in PubMed were added by referring to several PC compound review papers (13, 14, 15).

Finally, terpenes included in the herbal compound databases TCMSP (19), TCMID (20), and TM-MC (21) were added. The collected PC-terpenes were classified as lanostane-type triterpenes, seco-lanostane-type triterpenes, pentacyclic triterpenes, tricyclic diterpenes, sterols, and other classes using the ChEBI database (22) and manual curation.

For the collected PC terpenes, InChiKey and 3D structural information were searched using the PubChemPy library and manually curated; 125 compounds whose 3D molecular structures were retrieved from the PubChem Database (23) were selected as druggable compounds.

#### Method of Collecting Human-derived Druggable Proteomes

Druggable proteome information was collected from the Human Protein Atlas (HPA) (24). Druggable compounds primarily act on human protein targets such as enzymes, transporters, ion channels, and receptors. The proteome list consisted of 812 proteins categorized into enzymes, transporters, ion channels, and receptors known to interact with FDA-approved drugs provided by the HPA.

Structural information on human-derived proteins was obtained from the AlphaFold (AF) database (25). AF, a database that provides artificial-intelligence-based protein structure prediction results, also provides structural prediction information for human-derived proteins. After downloading the human-derived protein information of AF, 869 proteins included in the HPA druggable proteome list were extracted and selected as druggable proteomes for molecular docking.

#### Large-scale Molecular Docking Method using Terpenes and Druggable Proteomes

Druggable compounds and proteomes were transformed into pdbqt form, a structure that can be used for MDA, using OpenBabel software (26). Large-scale MDA was calculated by considering all cases in which the selected compounds and proteins were paired (125 compounds × 869 proteomes = 108,625 pairs). MDA was performed using Python and AutoDock vina software (27). The MDA parameters were set as follows: exhaustiveness, maximum value of 100; box size, maximum value [126, 126, 126].

In the compound-protein pair derived from the MDA results, 100 proteins with the lowest binding affinities for each of the 125 druggable compounds were selected as the main dockable proteomes for each compound. To identify the proteins that interact with each terpene class, we calculated the probability that dockable proteomes interact with each terpene class.

#### Over-representation analysis method using unique compounds of Poria cocos

ORA was performed using lanostane and seco-lanostane triterpenes, which are unique PC compounds. The 125 components subjected to the docking analysis were classified into lanostane, seco-lanostane, and non-lanostane groups. Interactions with terpenes were confirmed for each protein, and the number of terpenes with which the proteins interacted in each terpene group was counted. The number of proteins counted was expressed as a probability by dividing the total number of proteins in each terpene group. Subsequently, proteins with a docking probability of 50% or more in the lanostane and seco-lanostane groups, and a docking probability of 50% or less in the non-lanostane group were defined as proteins interacting with PC-specific terpenes. ORA was performed using the EnrichR platform for proteins interacting with PC-specific terpenes (28). For ORA, the KEGG (29), GO biological process (30), and OMIM disease (31) databases were used, and the results with the top 10 combined scores were selected as the ORA results of the unique compounds of PC.

#### Over-representation Analysis Method using Main Dockable Proteomes

ORA was performed using the main dockable proteome of the 125 components. ORA used the Enrichr API (28) provided by Python gseapy library (32) and performed analysis using the ‘KEGG_2021_Human’ gene set (29). After ORA, pathways that met the criteria of the Benjamini-Hochberg (BH) procedure (p<0.0001) for all 125 druggable compounds were selected as statistically significant (33). The frequency of all pathways found in the significant pathways of each component was summed, and the summed value was defined as the degree of significant MOA of the PC-terpenes.

However, this analysis could not eliminate selection bias arising from the druggable proteome list. Relative statistical analysis was performed using a permutation test to remove selection bias (34). A random gene set was created by randomly extracting 100 proteomes from the 869 druggable proteomes. A permutation list was constructed by grouping 125 random gene sets, and 1000 permutation lists were constructed. From the list of 1000 permutations, the same method used for docking-derived proteins was repeated to extract 1000 significant random pathways (125 random gene sets × 1000 permutations = 125,000 ORA scores). The number of occurrences of each significant pathway derived from docking was counted, and the same process was repeated for 1000 permutation sets. The sum of the significant pathways in the druggable proteomes was compared with the sum of the significant pathways in the list of 1000 permutations. The rank of the sum of the significant pathways in the druggable proteomes among all the permutated significant pathways was the p-value. Fifty-four pathways for which the number of significant pathways appeared within the top 5% (p<0.05) were selected as statistically significant final pathways.

### Method for producing *Poria Cocos*-induced transcriptomes in SW1783 and HT29 cell lines

#### Chemicals and reagents

Dulbecco’s modified Eagle’s medium (DMEM), phosphate-buffered saline (PBS), TrypLE Express, penicillin-streptomycin, and fetal bovine serum (FBS) were purchased from Gibco (Grand Island, NY, USA). Leibovitz’s L-15 medium was purchased from the American Type Culture Collection (ATCC, Manassas, VA, USA). The cell culture flasks and multiwell culture plates were purchased from Thermo Fisher Scientific (Waltham, MA, USA). Dimethyl sulfoxide (DMSO), wortmannin, LY294002, Thioridazine were purchased from Sigma-Aldrich (St. Louis, MO, USA), and QIAzol lysis reagents were purchased from Qiagen (Germantown, MD, USA). The Ez-Cytox Cell Viability Assay Kit was purchased from Dogen Bio (Seoul, Korea).

#### Preparation of hot water extract of PC

Dried *Poria cocos* Wolf (Polyporaceae) were supplied by Kwangmyung-dang Medicinal Herbs Co. (Ulsan, Republic of Korea), and its morphology was carefully validated by Dr. Goya Choi, Herbal Medicine Resources Research Center, Korea Institute of Oriental Medicine (KIOM), Republic of Korea. A voucher specimen (2-22-0221) was deposited in the Korean Herbarium of the Standard Herbal Resources of KIOM (Naju, Republic of Korea). The crushed material was extracted at 100 ± 3L for 3 h using water refluxed system (MS-DM609, Misung Scientific, Yangu, Republic of Korea) and filtered through 5-µm cartridge filter (KOC Biotech, Daejeon, Republic of Korea). The filtrates were concentrated under 40L using a rotary evaporator (Ev-1020t, SciLab, Seoul, Republic of Korea), and then freeze dried (LP20, IlShin Biobase, Dongduchen, Republic of Korea). The yield of PC was 24.2 %, and final extracts were homogenized and stored in an airtight container in a 4L cold room. For the *in vitro* investigation, homogenous PC was vigorously vortexed for 30 min at room temperature in phosphate-buffered saline (PBS; Thermo Fisher Scientific, Rockford, IL, USA) containing 2% DMSO and then sterilized by filtering through a 0.22-µm membrane. The stock solution of PC (10 mg/mL) was aliquoted in a 1.5-mL tube and stored at −80°C until use.

#### Cell culture

Human astrocytoma cell line SW1783 (HTB-13) and human colorectal adenocarcinoma cell line HT29 (HTB-38) were purchased from American Type Culture Collection (ATCC, Manassas, VA, USA). SW1783 cells were maintained in Leibovitz’s L-15 medium supplemented with 10% (v/v) heat-inactivated FBS, 100 IU/mL penicillin, and 100 mg/mL streptomycin at 37 °C, without CO_2_ incubator. HT29 cells were maintained in DMEM medium supplemented with 10% (v/v) heat-inactivated FBS, 100 IU/mL penicillin, and 100 mg/mL streptomycin at 37 °C, 5% CO_2_ incubator. The cells were sub-cultured every three or four days, depending on the cell density.

#### Drug treatment

One day before drug treatment, SW1783 and HT29 cells were plated at 1.5 × 10^5^ or 5 × 10^5^ cells/well, respectively in a 6-well plate containing 3 mL growth medium. The cells were exposed to 20 (Low), 100 (Middle), 500 (High) µg/mL by treatment with 150 µL per well of 0.4, 2, 10 mg/mL PC. There was no cytotoxicity at a high dose (500 µg/mL), which was confirmed using an Ez-Cytox cell viability assay kit. PBS with 2% DMSO was used as the vehicle and wortmannin, LY294002, thioridazine were treated at a concentration of 10 µM as a positive control. After 24 h of drug treatment, the cells were washed three times with ice-cold PBS. The total cell lysate was prepared with add QIAzol lysis reagent and stored in a −70 °C deep freezer until RNA extraction.

#### RNA preparation for RNA-seq

SW1783 and HT29 cells were subjected to total RNA extraction using the QIAzol lysis reagent according to the manufacturer’s instructions. The concentration of isolated RNA was determined using an Agilent RNA 6000 Nano Kit (Agilent Technologies, Waldbronn, Germany). RNA concentration was determined using a QuantitTM RiboGreen RNA assay kit (R11490, Thermo Fisher Scientific), and RNA quality was assessed by determining the RNA integrity number (>7) and 28S:18S ribosomal RNA ratio (>1.0) using an Agilent 2100 Bioanalyzer system (Agilent Technologies, Waldbronn, Germany). Total RNA was processed to prepare an mRNA sequencing library using the MGIEasy RNA Directional Library Prep kit (#1000006386; MGI Tech Co., Ltd., China) according to the manufacturer’s instructions. The library was quantified using the QauntiFluor® ssDNA System (E3190, Promega Corporation, WI, USA). The prepared DNA nanoballs were sequenced on an MGISeq system (MGI Tech Co., Ltd., China) with 100 bp paired-end reads.

#### RNA-seq preprocessing and differential gene expression analysis

The quality of raw RNA-seq data was evaluated using FastQC (v0.11.9). Adapter sequences were removed from the reads using TrimGalore (v0.6.6). The cleaned reads were then aligned to the human reference genome GRCh38 (hg19) using STAR (v2.7.9 a) with default parameters. The transcript abundance per gene was quantified using RSEM (v1.3.3) with the gene annotation GRCh38.84. The expected read counts and transcripts per million (TPM) were used as the gene expression levels for further analyses. Differential gene expression analysis between the treatment and vehicle groups was conducted using the Wald test implemented in the R package DESeq2 (v1.38.2). Wald statistic was used to rank genes, and this ranked list of genes was employed for gene set enrichment analysis (GSEA). The aforementioned analysis of RNA-seq data was performed using R (v4.2.2, R Foundation for Statistical Computing, Vienna, Austria). The RNA sequence data were deposited in the Gene Expression Omnibus with accession number (XXX).

### Gene Set Enrichment Analysis and Cluster Analysis of Pathways

GSEA was performed using the DTA results obtained from the cell lines (35). GSEA was performed for curated KEGG pathway gene sets in the Molecular Signature Database (MSigDB v7.5.1) (36) using the fgsea package (v1.24) (37) in R (v4.2.2) with parameters of minimum size 15, maximum size 500, and 100,000 permutations. Statistical significance of the GSEA results was evaluated by adjusting the p-value using the BH procedure. GSEA results were visualized as a heatmap using Pheatmap (v.1.0.12), and only pathways with results satisfying at least one statistical significance in the three PC concentrations were included in the heatmap. A pathway heatmap was used for hierarchical clustering analysis using correlation distance and complete methods (38).

### Diffusion Network Construction

#### Method for Construction of Alzheimer’s diffusion network based on disease pathway

A diffusion network (DN) was constructed to explain the connection between the MDA and DTA results. DN was created using the KAGS, which is a significant pathway in both MDA ORA and DTA GSEA results. Starting from the MDA results, a new layer was created with proteins exceeding the score (n≥950) based on the protein-protein interaction (PPI) combined score provided by the string database (39). The endpoints of the network used differentially expressed genes (DEGs) derived from DTA results.

For the MDA-based proteins used in the network, all proteins predicted to act on the AD pathway in the ORA analysis were collected from the 125 compounds used for docking. Subsequently, 14 proteins with an appearance frequency of 20% or more were selected as proteins based on MDA (n≥25). The DEGs used in the network were found to have a significant effect on the AD pathway, and the results for the medium concentration were used. Twenty-two proteins with an expression value greater than log_2_(0.5) or less than - log_2_(0.5) were selected as DEGs with an adjusted p-value<0.05 in the middle concentration. The protein set for the layer construction was constructed by downloading human protein information from the STRING database.

The network consists of five layers (Figure 2). The first and fifth layers comprised MDA-based proteins and DEGs, respectively. In the protein set, excluding the proteins included in the first and fifth layers, those with a PPI combined score of 950 or higher were selected as the proteins of the second layer. In the first layer and the second layer, a network was built by connecting interactions with a PPI score of 950 or higher. The third and fourth layers were constructed in the same way as the second layer, but all proteins that appeared once in the network were excluded from layer construction. The basic shape of the diffusion network was completed by connecting interactions with PPI scores of 950 or higher between proteins belonging to each protein set in the fourth and fifth layers (Step 1).

The network was then updated using the backpropagation method. When the fourth and fifth layers were connected, not all proteins belonging to the fourth layer were connected to the fifth layer. All the nodes in the fourth layer were deleted, except those that had at least one connection with the fifth layer. Similarly, the diffusion network was completed by sequentially deleting the protein nodes of the third, second, and first layers (Step 2). Network construction was performed using Python and network visualization was performed using Cytoscape software (v3.9.1) (40).

#### Submodule Construction Method for Identification of Modes of Action

To confirm the pathological MOA of Alzheimer’s disease in the constructed DN, a new DN was reconstructed using KAGS. First, proteins belonging to the KAGS were color-mapped onto the ADN to determine how many AD-related genes were included in the network. Next, in the ADN, the network was reconstructed using only the proteins included in KAGS. All nodes, except for proteins included in the KAGS, were removed from the ADN. Subsequently, the nodes with no edges were removed. The reconstructed network was relocated for each layer. The second layer was rearranged into five groups based on its connection to the same proteins belonging to the first layer. The third layer was also divided into five groups in a similar manner. The third layer groups were used to identify the MOA points of PC in the KEGG pathways and KEGG mappers. The group that acted on the submodule in the pathway was connected by adding submodule nodes. All the above visualizations were performed using Cytoscape software (version 3.9.1).

## Data availability statement

The raw sequence and processed data were deposited in the NCBI Gene Expression Omnibus (GEO, https://www.ncbi.nlm.nih.gov/geo/) with accession number GSE232862 and GSE232868.

## Acknowledgments

This research was funded by the research program of the Korea Institute of Oriental Medicine [grant number KSN1731122].

## Author contributions

M.S, P. contributed to the research design, analyzed data, and drafted the manuscript; S.Y, L and H.S, L. drafted the manuscript; J.M, Y conducted data design and production and drafted the manuscript. All authors have read and agreed to the published version of the manuscript.

## Reference

1. Breitmaier E. Terpenes: flavors, fragrances, pharmaca, pheromones: John Wiley & Sons; 2006.

2. Wang WH, Dong HJ, Yan RY, Li H, Li PY, Chen P, et al. Comparative study of lanostane-type triterpene acids in different parts of Poria cocos (Schw.) Wolf by UHPLC-Fourier transform MS and UHPLC-triple quadruple MS. J Pharmaceut Biomed. 2015;102:203–14.

3. Nukaya H, Yamashiro H, Fukazawa H, Ishida H, Tsuji K. Isolation of inhibitors of TPA-induced mouse ear edema from hoelen, Poria cocos. Chemical and pharmaceutical bulletin. 1996;44(4):847–9.

4. Jeong M-j, Kang K-w, Kang J-y, Yoon J-h, Choi Y-m, Kim H-j, et al. An Overview of the Applicability of Oryung-san as an Antihypertensive Agent. The Journal of Internal Korean Medicine. 2017;38(4):443–54.

5. Lee JH, Kim JI, Baeg MK, Sunwoo YY, Do K, Lee JH, et al. Effect of Samryungbaekchul-san Combined with Otilonium Bromide on Diarrhea-Predominant Irritable Bowel Syndrome: A Pilot Randomized Controlled Trial. J Clin Med. 2019;8(10).

6. Ko SJ, Park JW, Lee JH, Cho SH, Lee J, Nam S, et al. Herbal medicine Yukgunja-tang for functional dyspepsia protocol for a systematic review of randomized controlled trials. Medicine. 2018;97(40).

7. Li H, Li Q, Yang X, Wang Y, Tian X, Chen X, et al. Clinical study on Gui Pi Tang treating the depression and improving the life quality in elderly patients. China Journal of Traditional Chinese Medicine and Pharmacy. 2014;29(6):1855–9.

8. Rios JL. Chemical Constituents and Pharmacological Properties of Poria cocos. Planta Med. 2011;77(7):681–91.

9. Chang R. Functional properties of edible mushrooms. Nutr Rev. 1996;54(11 Pt 2):S91-3.

10. Bemis GW, Murcko MA. The properties of known drugs. 1. Molecular frameworks. J Med Chem. 1996;39(15):2887-93.

11. Hu Y, Stumpfe D, Bajorath J. Computational Exploration of Molecular Scaffolds in Medicinal Chemistry. Journal of Medicinal Chemistry. 2016;59(9):4062–76.

12. Park M, Baek S-J, Park S-M, Yi J-M, Cha S. Comparative study of the mechanism of natural compounds with similar structures using docking and transcriptome data for improving in silico herbal medicine experimentations. bioRxiv. 2023:2023.04.23.538005.

13. Wang YZ, Zhang J, Zhao YL, Li T, Shen T, Li JQ, et al. Mycology, cultivation, traditional uses, phytochemistry and pharmacology of Wolfiporia cocos (Schwein.) Ryvarden et Gilb.: a review. J Ethnopharmacol. 2013;147(2):265–76.

14. Lu J, Tian J, Zhou L, Meng LJ, Chen ST, Ma CY, et al. Phytochemistry and Biological Activities of Poria. J Chem-Ny. 2021;2021.

15. Nie A, Chao Y, Zhang X, Jia W, Zhou Z, Zhu C. Phytochemistry and Pharmacological Activities of Wolfiporia cocos (F.A. Wolf) Ryvarden & Gilb. Front Pharmacol. 2020;11:505249.

16. Fan J, Fu A, Zhang L. Progress in molecular docking. Quantitative Biology. 2019;7:83–9.

17. Dobin A, Gingeras TR. Mapping RNA-seq Reads with STAR. Curr Protoc Bioinformatics. 2015;51:11 4 1-4 9.

18. Chen L, Zhang YH, Zhang Z, Huang T, Cai YD. Inferring Novel Tumor Suppressor Genes with a Protein-Protein Interaction Network and Network Diffusion Algorithms. Mol Ther Methods Clin Dev. 2018;10:57–67.

19. Ru JL, Li P, Wang JN, Zhou W, Li BH, Huang C, et al. TCMSP: a database of systems pharmacology for drug discovery from herbal medicines. J Cheminformatics. 2014;6.

20. Huang L, Xie D, Yu Y, Liu H, Shi Y, Shi T, et al. TCMID 2.0: a comprehensive resource for TCM. Nucleic Acids Res. 2018;46(D1):D1117–D20.

21. Kim SK, Nam S, Jang H, Kim A, Lee JJ. TM-MC: a database of medicinal materials and chemical compounds in Northeast Asian traditional medicine. BMC Complement Altern Med. 2015;15:218.

22. Degtyarenko K, de Matos P, Ennis M, Hastings J, Zbinden M, McNaught A, et al. ChEBI: a database and ontology for chemical entities of biological interest. Nucleic Acids Res. 2008;36(Database issue):D344-50.

23. Kim S, Thiessen PA, Bolton EE, Chen J, Fu G, Gindulyte A, et al. PubChem Substance and Compound databases. Nucleic Acids Res. 2016;44(D1):D1202–13.

24. Uhlen M, Bjorling E, Agaton C, Szigyarto CA, Amini B, Andersen E, et al. A human protein atlas for normal and cancer tissues based on antibody proteomics. Mol Cell Proteomics. 2005;4(12):1920–32.

25. Jumper J, Evans R, Pritzel A, Green T, Figurnov M, Ronneberger O, et al. Highly accurate protein structure prediction with AlphaFold. Nature. 2021;596(7873):583-9.

26. O’Boyle NM, Banck M, James CA, Morley C, Vandermeersch T, Hutchison GR. Open Babel: An open chemical toolbox. J Cheminform. 2011;3:33.

27. Trott O, Olson AJ. AutoDock Vina: improving the speed and accuracy of docking with a new scoring function, efficient optimization, and multithreading. J Comput Chem. 2010;31(2):455–61.

28. Chen EY, Tan CM, Kou Y, Duan Q, Wang Z, Meirelles GV, et al. Enrichr: interactive and collaborative HTML5 gene list enrichment analysis tool. BMC Bioinformatics. 2013;14:128.

29. Ogata H, Goto S, Sato K, Fujibuchi W, Bono H, Kanehisa M. KEGG: Kyoto Encyclopedia of Genes and Genomes. Nucleic Acids Res. 1999;27(1):29–34.

30. Ashburner M, Ball CA, Blake JA, Botstein D, Butler H, Cherry JM, et al. Gene ontology: tool for the unification of biology. The Gene Ontology Consortium. Nat Genet. 2000;25(1):25–9.

31. Hamosh A, Scott AF, Amberger JS, Bocchini CA, McKusick VA. Online Mendelian Inheritance in Man (OMIM), a knowledgebase of human genes and genetic disorders. Nucleic Acids Res. 2005;33(Database issue):D514-7.

32. Fang Z, Liu X, Peltz G. GSEApy: a comprehensive package for performing gene set enrichment analysis in Python. Bioinformatics. 2023;39(1).

33. Benjamini Y, Hochberg Y. Controlling the false discovery rate: a practical and powerful approach to multiple testing. Journal of the Royal statistical society: series B (Methodological). 1995;57(1):289–300.

34. Welch WJ. Construction of permutation tests. Journal of the American Statistical Association. 1990;85(411):693–8.

35. Subramanian A, Tamayo P, Mootha VK, Mukherjee S, Ebert BL, Gillette MA, et al. Gene set enrichment analysis: A knowledge-based approach for interpreting genome-wide expression profiles. P Natl Acad Sci USA. 2005;102(43):15545–50.

36. Liberzon A, Subramanian A, Pinchback R, Thorvaldsdottir H, Tamayo P, Mesirov JP. Molecular signatures database (MSigDB) 3.0. Bioinformatics. 2011;27(12):1739–40.

37. Korotkevich G, Sukhov V, Budin N, Shpak B, Artyomov MN, Sergushichev A. Fast gene set enrichment analysis. BioRxiv. 2016:060012.

38. Murtagh F, Contreras P. Algorithms for hierarchical clustering: an overview. Wires Data Min Knowl. 2012;2(1):86–97.

39. von Mering C, Huynen M, Jaeggi D, Schmidt S, Bork P, Snel B. STRING: a database of predicted functional associations between proteins. Nucleic Acids Research. 2003;31(1):258–61.

40. Shannon P, Markiel A, Ozier O, Baliga NS, Wang JT, Ramage D, et al. Cytoscape: a software environment for integrated models of biomolecular interaction networks. Genome Res. 2003;13(11):2498–504.

41. Del Prado-Audelo ML, Cortes H, Caballero-Floran IH, Gonzalez-Torres M, Escutia-Guadarrama L, Bernal-Chavez SA, et al. Therapeutic Applications of Terpenes on Inflammatory Diseases. Front Pharmacol. 2021;12:704197.

42. Ramos CJ, Antonetti DA. The role of small GTPases and EPAC-Rap signaling in the regulation of the blood-brain and blood-retinal barriers. Tissue Barriers. 2017;5(3):e1339768.

43. Baulch JE, Acharya MM, Agrawal S, Apodaca LA, Monteiro C, Agrawal A. Immune and Inflammatory Determinants Underlying Alzheimer’s Disease Pathology. J Neuroimmune Pharmacol. 2020;15(4):852–62.

44. Raghavendra S, Rao SJA, Kumar V, Ramesh CK. Multiple ligand simultaneous docking (MLSD): A novel approach to study the effect of inhibitors on substrate binding to PPO. Comput Biol Chem. 2015;59:81–6.

45. Li TH, Hou CC, Chang CL, Yang WC. Anti-hyperglycemic properties of crude extract and triterpenes from Poria cocos. Evid Based Complement Alternat Med. 2011;2011.

46. Chao CL, Huang HW, Su MH, Lin HC, Wu WM. The Lanostane Triterpenoids in Poria cocos Play Beneficial Roles in Immunoregulatory Activity. Life (Basel). 2021;11(2).

47. He J, Yang Y, Zhang F, Li Y, Li X, Pu X, et al. Effects of Poria cocos extract on metabolic dysfunction-associated fatty liver disease via the FXR/PPARalpha-SREBPs pathway. Front Pharmacol. 2022;13:1007274.

48. Sun YF, Liu ZQ, Pi ZF, Song FR, Wu JL, Liu S. Poria cocos could ameliorate cognitive dysfunction in APP/PS1 mice by restoring imbalance of A beta production and clearance and gut microbiota dysbiosis. Phytother Res. 2021;35(5):2678–90.

49. Na S-S, Chong M-S, Woo J-H, Kwon Y-O, Lee MK, Oh K-W. Poria cocos ethanol extract and its active constituent, pachymic acid, modulate sleep architectures via activation of GABAA-ergic transmission in rats. Journal of Biomedical and Translational Research. 2015;16(3):84–92.

50. Jia XJ, Ma LS, Li P, Chen MW, He CW. Prospects of Poria cocos polysaccharides: Isolation process, structural features and bioactivities. Trends Food Sci Tech. 2016;54:52–62.

51. Yokoyama A, Natori S, Aoshima K. Distribution of tetracyclic triterpenoids of lanostane group and sterols in the higher fungi especially of the Polyporaceae and related families. Phytochemistry. 1975;14(2):487–97.

52. Akihisa T, Nakamura Y, Tokuda H, Uchiyama E, Suzuki T, Kimura Y, et al. Triterpene acid from Poria cocos and their anti-tumor-promoting effects. J Nat Prod. 2007;70(6):948–53.

53. Tai T, Akahori A, Shingu T. Triterpenes of Poria cocos. Phytochemistry. 1993;32(5):1239–44.

54. Huang HT, Wang SL, Nguyen VB, Kuo YH. Isolation and Identification of Potent Antidiabetic Compounds from Antrodia cinnamomea-An Edible Taiwanese Mushroom. Molecules. 2018;23(11).

55. Zhu LX, Xu J, Zhang SJ, Wang RJ, Huang Q, Chen HB, et al. Qualitatively and quantitatively comparing secondary metabolites in three medicinal parts derived from Poria cocos (Schw.) Wolf using UHPLC-QTOF-MS/MS-based chemical profiling. J Pharmaceut Biomed. 2018;150:278–86.

56. Feng GF, Zheng Y, Sun YF, Liu S, Pi ZF, Song FR, et al. A targeted strategy for analyzing untargeted mass spectral data to identify lanostane-type triterpene acids in Poria cocos by integrating a scientific information system and liquid chromatography-tandem mass spectrometry combined with ion mobility spectrometry. Anal Chim Acta. 2018;1033:87–99.

57. Ling Y, Chen MC, Wang K, Sun ZL, Li ZX, Wu B, et al. Systematic screening and characterization of the major bioactive components of Poria cocos and their metabolites in rats by LC-ESI-MS(n). Biomed Chromatogr. 2012;26(9):1109–17.

58. Zhang GH, Wang HX, Xie WY, Wang Q, Wang X, Wang CY, et al. Comparison of triterpene compounds of four botanical parts from Poria cocos (Schw.) wolf using simultaneous qualitative and quantitative method and metabolomics approach. Food Res Int. 2019;121:666–77.

59. Ru J, Li P, Wang J, Zhou W, Li B, Huang C, et al. TCMSP: a database of systems pharmacology for drug discovery from herbal medicines. J Cheminform. 2014;6:13.

60. Liya W, Huijie W. Studies on the chemical constituents of Fuling (Poria cocos). Zhong cao yao= Chinese Traditional and Herbal Drugs. 1998;29(3):145–8.

61. Zou YT, Long F, Wu CY, Zhou J, Zhang W, Xu JD, et al. A dereplication strategy for identifying triterpene acid analogues in Poria cocos by comparing predicted and acquired UPLC-ESI-QTOF-MS/MS data. Phytochem Anal. 2019;30(3):292–310.

62. Wu LF, Wang KF, Mao X, Liang WY, Chen WJ, Li S, et al. Screening and Analysis of the Potential Bioactive Components of Poria cocos (Schw.) Wolf by HPLC and HPLC-MS(n) with the Aid of Chemometrics. Molecules. 2016;21(2).

63. Tai T, Shingu T, Kikuchi T, Tezuka Y, Akahori A. Isolation of lanostane-type triterpene acids having an acetoxyl group from sclerotia of Poria cocos. Phytochemistry. 1995;40(1):225–31.

64. Yasukawa K, Kaminaga T, Kitanaka S, Tai T, Nunoura Y, Natori S, et al. 3 beta-p-hydroxybenzoyldehydrotumulosic acid from Poria cocos, and its anti-inflammatory effect. Phytochemistry. 1998;48(8):1357–60.

65. Li G, Xu ML, Lee CS, Woo MH, Chang HW, Son JK. Cytotoxicity and DNA topoisomerases inhibitory activity of constituents from the sclerotium of Poria cocos. Arch Pharm Res. 2004;27(8):829–33.

66. Zhou L, Zhang YC, Gapter LA, Ling H, Agarwal R, Ng KY. Cytotoxic and anti-oxidant activities of lanostane-type triterpenes isolated from Poria cocos. Chem Pharm Bull. 2008;56(10):1459–62.

67. Zheng Y, Yang X-W. Two new lanostane triterpenoids from Poria cocos. J Asian Nat Prod Res. 2008;10(4):289–92.

68. Zheng Y, Yang XW. Poriacosones A and B: two new lanostane triterpenoids from Poria cocos. J Asian Nat Prod Res. 2008;10(7):640–6.

69. Cai TG, Cai Y. Triterpenes from the fungus Poria cocos and their inhibitory activity on nitric oxide production in mouse macrophages via blockade of activating protein-1 pathway. Chem Biodivers. 2011;8(11):2135–43.

70. Kanematsu A, Natori S. Triterpenoids of Hoelen (fuling), sclerotia of Poria cocos (Schw.) Wolf. I. Pharm Soc Jap J. 1970.

71. Fu M, Wang L, Wang XY, Deng BX, Hu X, Zou J. Determination of the Five Main Terpenoids in Different Tissues of Wolfiporia cocos. Molecules. 2018;23(8).

72. Zheng Y, Liu S, Xing JP, Zheng Z, Pi ZF, Song FR, et al. Equivalently Quantitative Ion Strategy with Quaternary Ammonium Cation Derivatization for Highly Sensitive Quantification of Lanostane-Type Triterpene Acids without Standards by Ultrahigh-Performance Liquid Chromatography-Tandem Mass Spectrometry (UHPLC-MS/MS). Anal Chem. 2018;90(23):13946–52.

73. Zan JF, Shen CJ, Zhang LP, Liu YW. Effect of Poria cocos hydroethanolic extract on treating adriamycin-induced rat model of nephrotic syndrome. Chin J Integr Med. 2017;23(12):916–22.

74. Tian S-S, Liu X-Q, Feng W-H, Zhang Q-W, Yan L-H, Wang Z-M, et al. Quality evaluation of Poria based on specific chromatogram and quantitative analysis of multicomponents. Zhongguo Zhong yao za zhi= Zhongguo Zhongyao Zazhi= China Journal of Chinese Materia Medica. 2019;44(7):1371–80.

75. Giner-Larza EM, Máñez S, Giner-Pons RM, Recio MC, Rıos J-L. On the anti-inflammatory and anti-phospholipase A2 activity of extracts from lanostane-rich species. J Ethnopharmacol. 2000;73(1-2):61–9.

76. Zheng Y, Yang X-W. Absorption of triterpenoid compounds from Indian bread (Poria cocos) across human intestinal epithelial (Caco-2) cells in vitro. Zhongguo Zhong yao za zhi= Zhongguo Zhongyao Zazhi= China Journal of Chinese Materia Medica. 2008;33(13):1596–601.

77. Akihisa T, Uchiyama E, Kikuchi T, Tokuda H, Suzuki T, Kimura Y. Anti-tumor-promoting effects of 25-methoxyporicoic acid A and other triterpene acids from Poria cocos. J Nat Prod. 2009;72(10):1786–92.

78. Cheng SJ, Castillo V, Sliva D. CDC20 associated with cancer metastasis and novel mushroom-derived CDC20 inhibitors with antimetastatic activity. Int J Oncol. 2019;54(6):2250–6.

79. Zhang Y, Hu G-S, Han Z-F, Xiao W, Wang Z-D, Bi Y-A, et al. Dynamic accumulation of three main triterpenic acids in submerged cultivation mycelium of Poria cocos. Zhongguo Zhong yao za zhi= Zhongguo Zhongyao Zazhi= China Journal of Chinese Materia Medica. 2013;38(9):1355–9.

80. Tai T, Shingu T, Kikuchi T, Tezuka Y, Akahori A. Triterpenes from the surface layer of Poria cocos. Phytochemistry. 1995;39(5):1165–9.

81. Wu YJ, Li S, Li HX, Zhao CZ, Ma H, Zhao XN, et al. Effect of a polysaccharide from Poria cocos on humoral response in mice immunized by H1N1 influenza and HBsAg vaccines. Int J Biol Macromol. 2016;91:248–57.

82. Zeng HL, Liu Q, Yu JG, Jiang XY, Wu ZL, Wang ML, et al. One-step separation of nine structural analogues from Poria cocos (Schw.) Wolf. via tandem high-speed counter-current chromatography. J Chromatogr B. 2015;1004:10–6.

83. Xia B, Zhou Y, Tan HS, Ding LS, Xu HX. Advanced ultra-performance liquid chromatography-photodiode array-quadrupole time-of-flight mass spectrometric methods for simultaneous screening and quantification of triterpenoids in Poria cocos. Food Chem. 2014;152:237–44.

84. Kikuchi T, Uchiyama E, Ukiya M, Tabata K, Kimura Y, Suzuki T, et al. Cytotoxic and apoptosis-inducing activities of triterpene acids from Poria cocos. J Nat Prod. 2011;74(2):137–44.

85. Ukiya M, Akihisa T, Tokuda H, Hirano M, Oshikubo M, Nobukuni Y, et al. Inhibition of tumor-promoting effects by poricoic acids G and H and other lanostane-type triterpenes and cytotoxic activity of poricoic acids A and G from Poria cocos. J Nat Prod. 2002;65(4):462–5.

86. Chen BS, Zhang RJ, Han JJ, Zhao RL, Bao L, Huang Y, et al. Lanostane Triterpenoids with Glucose-Uptake-Stimulatory Activity from Peels of the Cultivated Edible Mushroom Wolfiporia cocos. J Agr Food Chem. 2019;67(26):7348–64.

87. Akihisa T, Mizushina Y, Ukiya M, Oshikubo M, Kondo S, Kimura Y, et al. Dehydrotrametenonic acid and dehydroeburiconic acid from Poria cocos and their inhibitory effects on eukaryotic DNA polymerase alpha and beta. Biosci Biotech Bioch. 2004;68(2):448–50.

88. Zhong Z, Liu J. Triterpenes isolated from Poria cocos by Derivazations. Zhong yao cai= Zhongyaocai= Journal of Chinese Medicinal Materials. 2002;25(4):247–50.

89. Yang CH, Zhang SF, Liu WY, Zhang ZJ, Liu JH. Two new triterpenes from the surface layer of Poria cocos. Helv Chim Acta. 2009;92(4):660–7.

90. Lee JH, Lee YJ, Shin JK, Nam JW, Nah SY, Kim SH, et al. Effects of triterpenoids from Poria cocos Wolf on the serotonin type 3A receptor-mediated ion current in Xenopus oocytes. Eur J Pharmacol. 2009;615(1-3):27–32.

91. Yang LB, Qin B, Feng SM, Liu SJ, Song XM. A new triterpenoid from traditional Chinese medicine Poria cocos. J Chem Res. 2010(10):553–4.

92. Li SN, Zhang JX, Li SL, Liu CM, Liu S, Liu ZQ. Extraction and separation of lactate dehydrogenase inhibitors from Poria cocos (Schw.) Wolf based on a hyphenated technique and in vitro methods. J Sep Sci. 2017;40(8):1773–83.

93. Lee SR, Lee S, Moon E, Park HJ, Park HB, Kim KH. Bioactivity-guided isolation of anti-inflammatory triterpenoids from the sclerotia of Poria cocos using LPS-stimulated Raw264.7 cells. Bioorg Chem. 2017;70:94–9.

94. Lv CX, Li Q, Zhang YW, Sui ZY, He BS, Xu HR, et al. A UFLC-MS/MS method with a switching ionization mode for simultaneous quantitation of polygalaxanthone III, four ginsenosides and tumulosic acid in rat plasma: application to a comparative pharmacokinetic study in normal and Alzheimer’s disease rats. J Mass Spectrom. 2013;48(8):904–13.

95. Yang P-F, Liu C, Wang H-Q, Li J-C, Wang Z-Z, Xiao W, et al. Chemical constituents of Poria cocos. Zhongguo Zhong yao za zhi= Zhongguo Zhongyao Zazhi= China Journal of Chinese Materia Medica. 2014;39(6):1030–3.

96. Ji B, Zhao YL, Yu PP, Yang B, Zhou C, Yu ZG. LC-ESI-MS/MS method for simultaneous determination of eleven bioactive compounds in rat plasma after oral administration of Ling-Gui-Zhu-Gan Decoction and its application to a pharmacokinetics study. Talanta. 2018;190:450–9.

97. Song ZH, Bi KS, Luo X, Chan K. The isolation, identification and determination of dehydrotumulosic acid in Poria cocos. Anal Sci. 2002;18(5):529–31.

98. Shingu T, Tai T, Akahori A. A lanostane triterpenoid from Poria cocos. Phytochemistry. 1992;31(7):2548–9.

99. Jin J, Zhou RR, Xie J, Ye HX, Liang XJ, Zhong C, et al. Insights into Triterpene Acids in Fermented Mycelia of Edible Fungus Poria cocos by a Comparative Study. Molecules. 2019;24(7).

100. Che S, Li Q, Huo Y-S, Chen X-H, Bi K-S. RP-HPLC simultaneous determination of five triterpenoid acids in different parts of Poria cocos by UV wavelengths switch. Yao xue xue bao= Acta Pharmaceutica Sinica. 2010;45(4):494–7.

101. Eom S, Kim YS, Lee SB, Noh S, Yeom HD, Bae H, et al. Molecular determinants of α3β4 nicotinic acetylcholine receptors inhibition by triterpenoids. Biological and Pharmaceutical Bulletin. 2018;41(1):65–72.

102. Lai KH, Lu MC, Du YC, El-Shazly M, Wu TY, Hsu YM, et al. Cytotoxic Lanostanoids from Poria cocos. J Nat Prod. 2016;79(11):2805–13.

103. Lee S, Lee S, Roh HS, Song SS, Ryoo R, Pang C, et al. Cytotoxic Constituents from the Sclerotia of Poria cocos against Human Lung Adenocarcinoma Cells by Inducing Mitochondrial Apoptosis. Cells-Basel. 2018;7(9).

104. Lee S, Choi E, Yang SM, Ryoo R, Moon E, Kim SH, et al. Bioactive compounds from sclerotia extract of Poria cocos that control adipocyte and osteoblast differentiation. Bioorg Chem. 2018;81:27–34.

105. Zhang N, Li Z, Li J, Liu J, Dai J, Li Y. Advances in the research of constituents and pharmacological effects of Poria cocos (Schw.) Wolf. World Sci Technol Modern Tradition Chin Med Mater Med. 2019;21:220–33.

106. Qian Q, Zhou N, Qi PC, Zhang YQ, Mu XY, Shi XW, et al. A UHPLC-QTOF-MS/MS method for the simultaneous determination of eight triterpene compounds from Poria cocos (Schw.) Wolf extract in rat plasma: Application to a comparative pharmacokinetic study. J Chromatogr B. 2018;1102:34–44.

107. Zhu LX, Xu J, Wang RJ, Li HX, Tan YZ, Chen HB, et al. Correlation between Quality and Geographical Origins of Poria cocos Revealed by Qualitative Fingerprint Profiling and Quantitative Determination of Triterpenoid Acids. Molecules. 2018;23(9).

108. Wang S, Jiang Y, Zhu N, Liu Y, Shi R, Yang X, et al. Determination and isolation of the chemical constituents of Poria cocos. Journal of Beijing University of Traditional Chinese Medicine. 2010;33(12):841–4.

109. Tai T, Akahori A, Shingu T. Triterpenoids from Poria cocos. Phytochemistry. 1991;30(8):2796–7.

110. Li S, Wang Z, Gu R, Zhao Y, Huang W, Wang Z, et al. A new epidioxy-tetracyclic triterpenoid from Poria cocos Wolf. Nat Prod Res. 2016;30(15):1712–7.

111. Chen T, Hua L, Chou GX, Mao XD, Zou XL. A Unique Naphthone Derivative and a Rare 4,5-seco-Lanostane Triterpenoid from Poria cocos. Molecules. 2018;23(10).

112. Chihara G, Hamuro J, Maeda Y, Arai Y, Fukuoka F. Antitumor polysaccharide derived chemically from natural glucan (pachyman). Nature. 1970;225(5236):943-4.

113. Feng GF, Li SZ, Liu S, Song FR, Pi ZF, Liu ZQ. Targeted Screening Approach to Systematically Identify the Absorbed Effect Substances of Poria cocos in Vivo Using Ultrahigh Performance Liquid Chromatography Tandem Mass Spectrometry. J Agr Food Chem. 2018;66(31):8319–27.

114. Lee S, Lee D, Lee SO, Ryu JY, Choi SZ, Kang KS, et al. Anti-inflammatory activity of the sclerotia of edible fungus, Poria cocos Wolf and their active lanostane triterpenoids. J Funct Foods. 2017;32:27–36.

115. Peng CH, Yang MY, Yang YS, Yu CC, Wang CJ. Antrodia cinnamomea Prevents Obesity, Dyslipidemia, and the Derived Fatty Liver via Regulating AMPK and SREBP Signaling. Am J Chinese Med. 2017;45(1):67–83.

116. Lee D, Lee S, Shim SH, Lee HJ, Choi Y, Jang TS, et al. Protective effect of lanostane triterpenoids from the sclerotia of Poria cocos Wolf against cisplatin-induced apoptosis in LLC-PK1 cells. Bioorg Med Chem Lett. 2017;27(13):2881–5.

117. Wang M, Chen DQ, Chen L, Liu D, Zhao H, Zhang ZH, et al. Novel RAS Inhibitors Poricoic Acid ZG and Poricoic Acid ZH Attenuate Renal Fibrosis via a Wnt/β-Catenin Pathway and Targeted Phosphorylation of smad3 Signaling. J Agr Food Chem. 2018;66(8):1828–42.

118. Chen L, Cao G, Wang M, Feng YL, Chen DQ, Vaziri ND, et al. The Matrix Metalloproteinase-13 Inhibitor Poricoic Acid ZI Ameliorates Renal Fibrosis by Mitigating Epithelial-Mesenchymal Transition. Mol Nutr Food Res. 2019;63(13).

119. Dong HJ, Wu PP, Yan RY, Xu QH, Li H, Zhang FB, et al. Enrichment and separation of antitumor triterpene acids from the epidermis of Poria cocos by pH-zone-refining counter-current chromatography and conventional high-speed counter-current chromatography. J Sep Sci. 2015;38(11):1977–82.

120. Jin Y, Zhang L, Chen L, Chen Y, Cheung PCK, Chen LG. Effect of culture media on the chemical and physical characteristics of polysaccharides isolated from Poria cocos mycelia. Carbohyd Res. 2003;338(14):1507–15.

121. Lin H-C, Song Y-Y, Huang Y-C, Chang W-L. A 4, 5-secolanostane triterpenoid from the sclerotium of Poria cocos. Journal of Medical Sciences. 2010;30(6):237-40.

122. Wang HL, Mukerabigwi JF, Zhang YN, Han L, Jiayinaguli T, Wang Q, et al. In vivo immunological activity of carboxymethylated-sulfated (1→3)-β-D-glucan from sclerotium of Poria cocos. Int J Biol Macromol. 2015;79:511–7.

123. Zhu LX, Xu J, Wu Y, Su LF, Lam KYC, Qi ER, et al. Comparative quality of the forms of decoction pieces evaluated by multidimensional chemical analysis and chemometrics: Poria cocos, a pilot study. J Food Drug Anal. 2019;27(3):766–77.

124. Zhang W, Cheng N, Wang Y, Zheng X, Zhao Y, Wang H, et al. Adjuvant activity of PCP-II, a polysaccharide from Poria cocos, on a whole killed rabies vaccine. Virus Res. 2019;270:197638.

125. Wang M, Chen DQ, Wang MC, Chen H, Chen L, Liu D, et al. Poricoic acid ZA, a novel RAS inhibitor, attenuates tubulo-interstitial fibrosis and podocyte injury by inhibiting TGF-β/Smad signaling pathway. Phytomedicine. 2017;36:243–53.

126. Xu S, Jiang W, Kuang Y, Wu X, Ma J, Jin P. Research advances on chemical constituents and bioactivities of Poria cocos (Schw.) Wolf. Northwest Pharmaceutical Journal. 2016;31(3):327–9.

127. Ding G, Wang Z-z, Zhang C-f, Sheng L-s. Study on HPLC fingerprint of the triterpene acids in Poria cocos. Zhongguo Zhong yao za zhi= Zhongguo Zhongyao Zazhi= China Journal of Chinese Materia Medica. 2002;27(10):756–8.

128. Li L, Yuan E, Gou N. Effects of water extract of poria on Helicobacter pylori inhibition and GES-1 cells proliferation. Modern Food Science & Technology. 2019;35(10):19–24.

129. Yang D, Chen Z, Liu Y, Zhao Y, Zhou J. Chemical constituents from Poria cocos (Schw.) Wolf. Anhui Agr Sci Bull. 2010;16(19):45–6.

130. Yu S-J, Tseng J. Fu-Ling, a Chinese herbal drug, modulates cytokine secretion by human peripheral blood monocytes. International journal of immunopharmacology. 1996;18(1):37–44.

