## Supplementary Figure 1 for "Effect of Terpenes from *Poria Cocos*: Verifying Modes of Action against Alzheimer’s disease Using Molecular Docking, Drug-induced Transcriptomes and Diffusion Network"

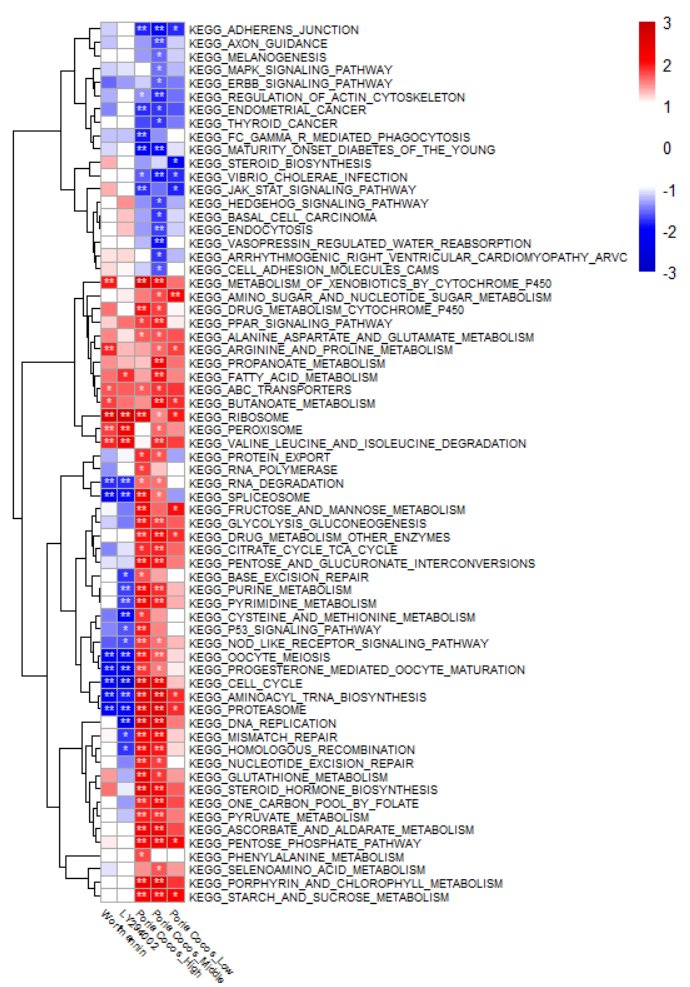

Supplementary Figure 1 GSEA results using the HT29 cell line-based PC-induced transcriptomes

GSEA was performed using the KEGG pathway gene set. Poria Cocos_High, Poria Cocos_Middle, and Poria Cocos_Low are the results of analyzing the gene expression values obtained by treating the PC water extract with high (500 ug/mL), medium (100 ug/mL), and low (20 ug/mL) concentrations, respectively. Wortmannin, LY294002, and Thioridazine are positive controls. Wortmannin and LY294002 are known to inhibit inflammation by acting on the cell cycle mechanism. GSEA, Gene set enrichment analysis. PC, Poria Cocos; KEGG, Kyoto Encyclopedia of Genes and Genomes.
