## Supplementary Figure 2 for "Effect of Terpenes from *Poria Cocos*: Verifying Modes of Action against Alzheimer’s disease Using Molecular Docking, Drug-induced Transcriptomes and Diffusion Network"

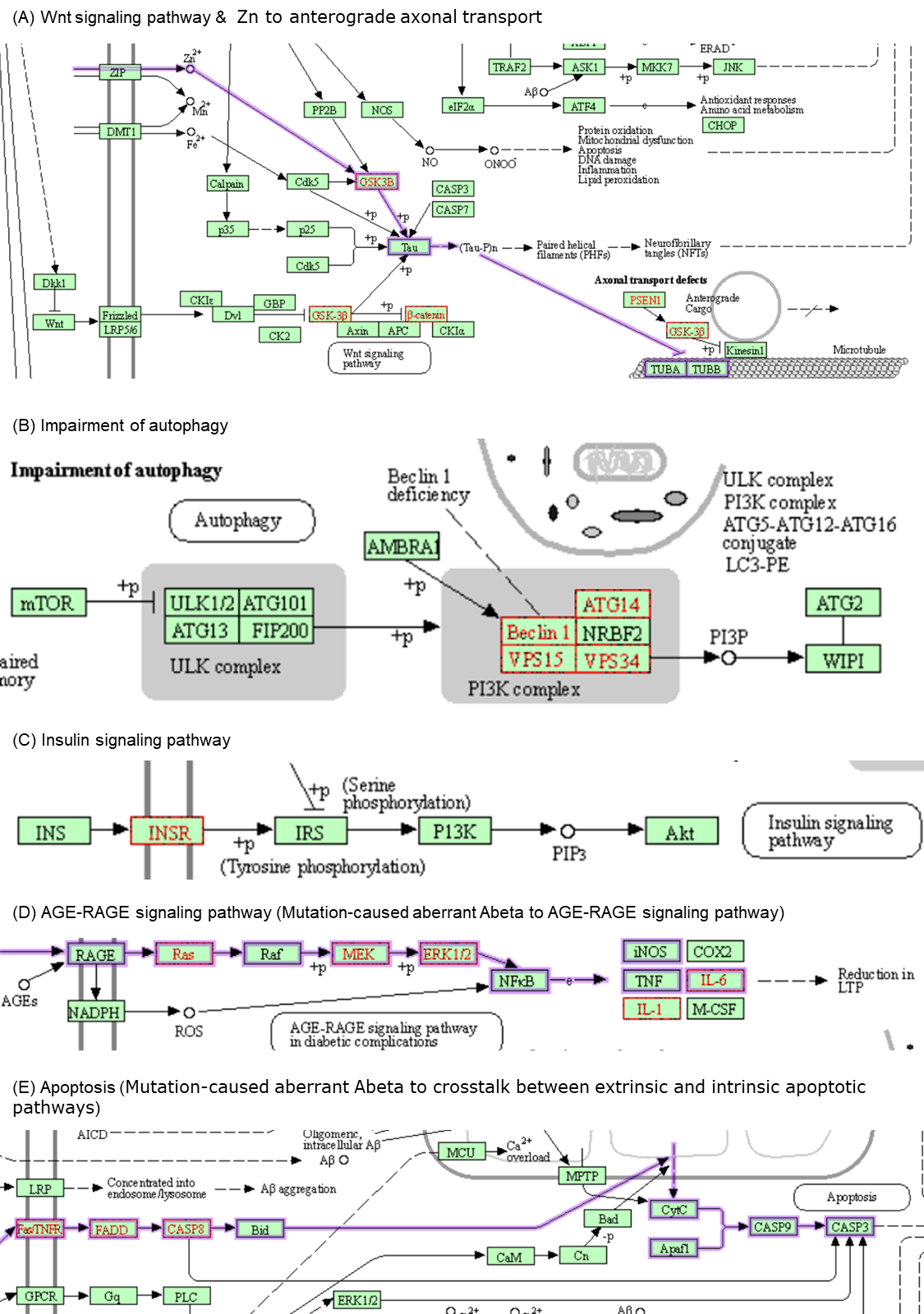

Supplementary Figure 2 Pathway of submodules of the KEGG AD pathway

Purple borders and lines represent submodules in the KEGG AD pathway. Red borders and letters are proteins whose expression values change in each submodule. KEGG, Kyoto Encyclopedia of Genes and Genomes; AD, Alzheimer's disease.
